# Tissue Engineered Elastic Cartilage-Mimetic Auricular Grafts for Ear Reconstruction

**DOI:** 10.1101/2025.10.27.684810

**Authors:** Philipp Fisch, Sandra Kessler, Simone Ponta, Anna Puiggalí-Jou, Guoliang Lyu, Killian Flégeau, Anastasiya Martyts, Florian Roth, David Fercher, Filippo M. Rijli, Daniel Simmen, Eva Novoa Olivares, Thomas Linder, Marcy Zenobi-Wong

## Abstract

Patients born with microtia, the congenital malformation of the external ear, face substantial psychosocial strain. Current reconstruction relies on harvesting rib cartilage, an invasive procedure associated with donor site morbidity and unnaturally stiff ears due to the use of hyaline cartilage. Tissue engineered auricles could overcome these drawbacks by providing patient-specific elastic cartilage without the need for rib harvest. Yet, key challenges such as fibrocartilage formation, inhomogeneous extracellular matrix formation and mechanical inferiority during *ex vivo* maturation remain, often leading to graft deformation and degradation *in vivo*. To address this gap, we integrated approaches maintaining the chondrogenic potential with growth factors, promoting elastic cartilage formation through stress-relaxing materials, and achieving homogeneous maturation by culturing grafts on an elevated bioreactor platform that enables uniform nutrient diffusion, using primary human auricular chondrocytes. Together these approaches resulted in the maturation of bioprinted auricular grafts that closely resemble native human auricular cartilage, demonstrated by the uniform distribution of elastin, glycosaminoglycans, and collagen II, and lack of collagen I. RNA sequencing revealed gene expression patterns consistent with the transition from fibrocartilage towards elastic cartilage. On a functional level, grafts achieved a compressive modulus of 1.1±0.03 MPa, matching that of native human auricular cartilage (1.0±0.1 MPa) and maintained their structural integrity for 6 weeks in a subcutaneous rat model, where they transitioned towards mature elastic fibers. These grafts represent the closest approximation of native elastic cartilage achieved *ex vivo* to date, bringing the field closer to a clinically viable, long-term therapy for children affected by microtia.

**One-sentence summary:** Bioprinted auricular grafts develop native-like elastic cartilage and advance towards a durable therapy for children with microtia.

## Introduction

Microtia, the congenital malformation of the external ear, represents a serious psychosocial burden for children and their parents, affecting about 1 to 4 children per 10’000 births (*1*). Children suffer from reduced self-esteem, increased anxiety, depression, and social isolation (*2–4*). As a rare or orphan disease and thus with limited interest of the biopharmaceutical industry, treatments for patients are limited (*5–8*).

To date, autologous costal cartilage reconstruction is the gold standard treatment for microtia patients (*9, 10*). However, since its development in the 1950s, progress in the treatment has been limited. Medical advances mainly focused on reducing the number of operative procedures, improving the ear framework sculpting and the cutaneous covering (*11–14*). The use of costal cartilage however remains vital for the procedure, often requiring patients to reach a specific age to have grown a sufficient amount of costal cartilage for harvest before surgery can be carried out (*15, 16*). This delay can increase the psychosocial distress of affected children (*17*). In addition, autologous costal cartilage reconstruction can lead to donor site morbidity including chest wall deformation, hypertrophic scarring and pain, and result in an impaired aesthetic outcome, extrusion of the cartilage framework, infections, hematoma, skin necrosis and framework resorption (*18, 19*).

Porous high-density polyethylene implants were developed in the 1980s to circumvent the harvest of rib cartilage (*20, 21*). However, these implants increased the complexity of the surgery with the need for a fascia flap and are associated with higher complication rates such as extrusion through the skin and infections (*22*). To avoid the implantation of an auricular framework, patients can further opt for osseointegrated epitheses (*23*). While these epitheses generally result in a better aesthetic outcome, they need to be replaced every 2-3 years, leading to high cost and are usually considered for failed auricular reconstructions (*24*). With the desire to obtain a real, natural ear rather than a synthetic one, patients prefer autologous costal cartilage reconstruction over polyethylene implants or epitheses (*2*).

Tissue engineering could substantially improve the treatment of microtia by manufacturing patient specific ears leading to an improved aesthetic outcome and omitting the harvest of costal cartilage (*25*). Such ears would allow to reduce the number of operative procedures and to perform the surgery at an earlier age, preventing children from the additional psychosocial strain by the delayed operation. Ultimately, tissue engineering could enable the replication of elastic cartilage resulting in a natural, real ear as opposed to the stiff costal cartilage frameworks used to date (*26, 27*).

Despite more than three decades of research (*28*) and numerous efforts to engineer auricular grafts for the treatment of microtia, no tissue engineered product has yet reached clinical translation. A major obstacle remains the dedifferentiation of chondrocytes during culture, leading to the formation of fibrocartilage rather than elastic cartilage *ex vivo* (*29, 30*). As a result, grafts are often implanted soon after fabrication. However, once exposed to physiological forces *in vivo*, insufficiently matured and mechanically weak grafts deform and lose their anatomical features (*25, 31*). To mitigate such deformations, three-dimensional printed external scaffolds made from polylactic acid (PLA) have been used to stabilize grafts during *in vivo* maturation (*32, 33*), but the inability to guide tissue maturation once implanted results in inhomogeneous extracellular matrix (ECM) deposition. The inhomogeneous ECM deposition in larger grafts further aggravates the deformations and loss of definition *in vivo* (*34–37*). In line with these challenges, auricular grafts previously developed in our group resulted in fibrocartilage (*38–40*) and ultimately failed *in vivo*, despite reaching 400 kPa, approximately one quarter of the stiffness of native auricular cartilage (Fig. S1).

To overcome these challenges, we developed strategies to mature auricular grafts with properties closely resembling native elastic cartilage (Fig. 1A-B). Using primary human auricular chondrocytes, we first show how the addition of growth factors during cell expansion (Fig. 1C), the incorporation of stress-relaxing properties in our bioink (Fig. 1D), and the adaptation of culture conditions (Fig. 1E) maintain the chondrogenic redifferentiation potential and improve tissue maturation. We then applied these strategies to bioprinted auricular grafts, matured on a bioreactor platform, and demonstrate their development into near-native auricular cartilage *ex vivo* (Fig. 1F). Finally, we assessed the structural stability of auricular grafts *in vivo* and lay the foundation for the generation of fully functional auricular grafts composed of mature elastic cartilage for the future treatment of children affected by microtia.

**Fig. 1.**
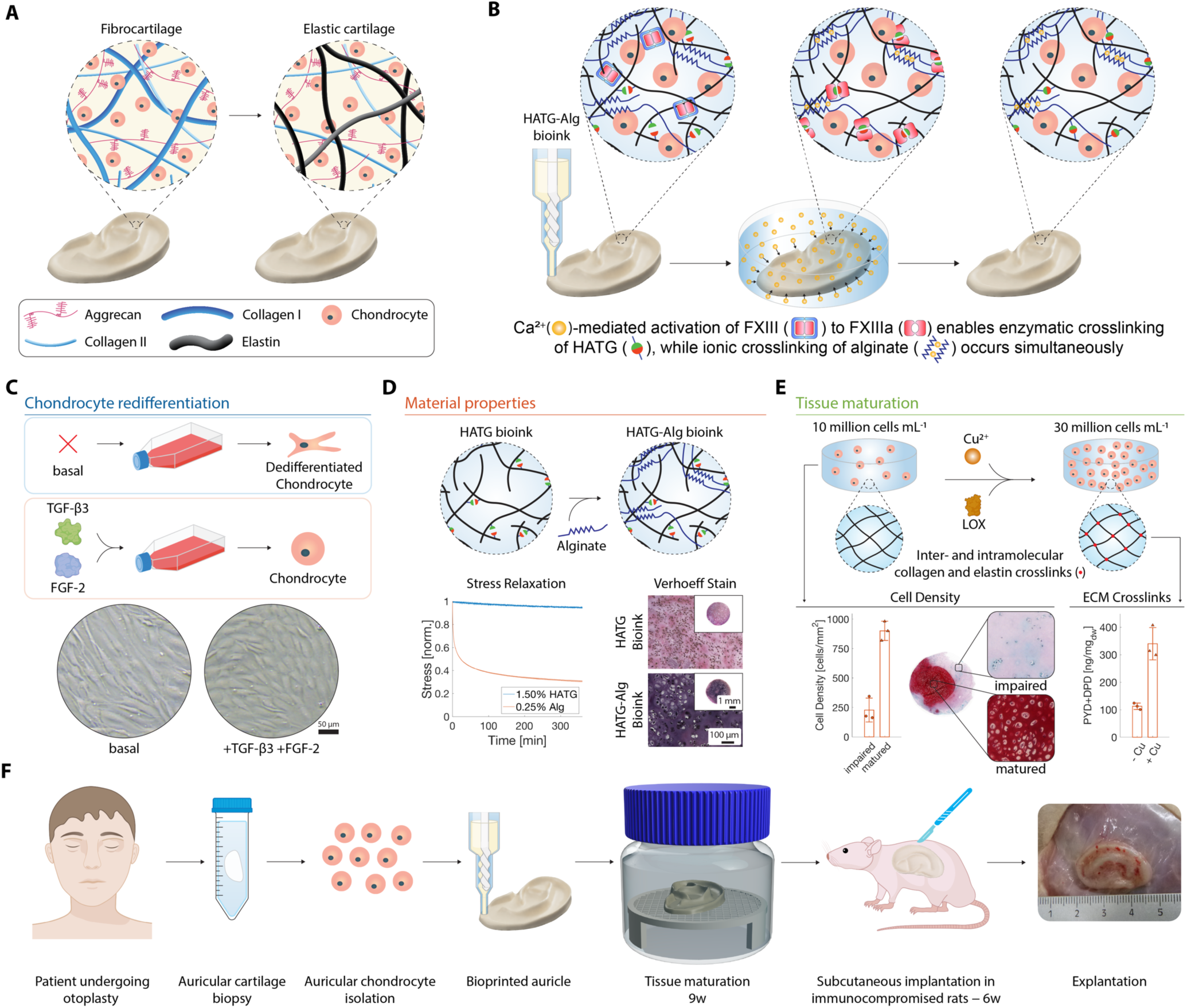
Towards tissue engineering elastic cartilage-mimetic auricular grafts. (**A**) Illustration of fibro- and elastic cartilage. (**B**) Bioink design combining calcium-triggered enzymatic crosslinking (CTEC) of hyaluronan transglutaminase (HATG) and ionic crosslinking of alginate (Alg). (**C-E**) Approaches to promote elastic cartilage formation. (**C**) Maintenance of the chondrogenic redifferentiation potential through the addition of growth factors such as FGF-2 and TGF-β3 during expansion and improved chondrocyte morphology. Scale bar: 50 μm. (**D**) Modification of the HATG bioink through the addition of alginate (HATG-Alg). In contrast to HATG hydrogels (1.5% HATG), alginate hydrogels (0.25% Alg) show distinct stress relaxation and HATG-Alg leads to the deposition of elastin (Verhoeff stain). (**E**) Improving tissue maturation by increasing cell density from 10 to 30 million cells per mL and through the addition of exogenous lysyl oxidase (LOX) and copper (Cu) to increase collagen and elastin crosslinking. Cell density in tissue engineered grafts in areas of matured and impaired tissue. Pyridinoline (PYD) and deoxypyridinoline (DPD) collagen crosslinks in grafts with (+Cu) or without (-Cu) copper supplementation. (**F**) *In vivo* evaluation of auricular grafts. Primary human auricular chondrocytes were isolated from auricular cartilage of otoplasty patients, bioprinted, cultured on an elevated bioreactor platform for 9 weeks and implanted subcutaneously in immunocompromised rats for 6 weeks.

## Results

### 2D expansion with FGF-2 and TGF-β3 improves auricular chondrocyte redifferentiation and prevents fibrocartilage formation

Chondrocytes are known to dedifferentiate during expansion, resulting in the formation of fibrocartilage (*41, 42*). Growth factors (GF) have been shown to perform a leading role in maintaining the chondrogenic redifferentiation potential during 2D expansion (*43, 44*). Fibroblast growth factor 2 (FGF-2) is particularly known for its leading role in the differentiation of neural crest cells towards a chondrogenic lineage, supporting role in cartilage formation (*45*) and maintaining the redifferentiation potential of chondrocytes during expansions (*43, 46*). The exact function of FGF-2 in auricular cartilage formation is however unclear. In addition, growth factors of the transforming growth factor β (TGF-β) family play an important role in cartilage homeostasis and ECM production and maintenance (*47*). Low doses of TGF-β further inhibit chondrocyte hypertrophy and terminal differentiation (*48*). We therefore investigated whether supplementing the media with FGF-2 and TGF-β3, as well as BMP-2 and BMP-9, during 2D expansion would enhance the redifferentiation potential of human auricular chondrocytes (hAUR, Fig. S2A).

Expanding hAUR in the presence of FGF-2 and TGF-β3 increased cell yield (Fig. S2B) and showed a trend towards higher expression of aggrecan, elastin and collagen I, while collagen II and X appeared unaffected (Fig. S2E-I). hAUR further showed a round morphology compared to a spread morphology under basal conditions (Fig. S2D,J) indicating the maintenance of their redifferentiation potential (*49*). While BMP-2 and BMP-9 impacted gene expression, neither increased elastin expression nor cell yield compared to hAUR expanded in the presence of FGF-2 alone.

We then combined hAUR expanded in the presence of GFs with our previously developed hyaluronan transglutaminase (HATG) bioink (*50*) and matured these grafts for 9 weeks (Fig. S3, Fig. S4). Conditions expanded with FGF-2 and TGF-β3 and supplemented with TGF-β3 during graft culture had the most pronounced effect on graft stiffness (Fig. S4A-C) and neocartilage formation (Fig. S4D), while no collagen I was observed at 9 weeks, indicating a shift away from fibrocartilage towards a more mature cartilage type. While BMP-2 and BMP-9 were able to prevent fibrocartilage formation, grafts were significantly softer than their TGF-β3 counterparts. Considering the risk of inducing chondrocyte hypertrophy using BMPs, we continued to focus on FGF-2 and TGF-β3. Despite the beneficial effects of FGF-2 and TGF-β3 on chondrocyte redifferentiation and shift away from fibrocartilage, grafts lacked elastin, a critical marker for elastic cartilage, indicating that additional factors may be necessary to induce elastic cartilage formation in grafts.

### Increased stress-relaxation through alginate supports graft maturation into elastic rather than fibrocartilage

Viscoelastic materials such as alginate (Alg) are known to promote chondrogenesis through mechanical confinement and stress relaxation, with faster stress relaxation promoting a chondrogenic phenotype and cartilaginous ECM deposition (*51, 52*). We therefore tested if supplementing Alg into our previously developed hyaluronan transglutaminase (HATG) bioink (*50*) would improve chondrogenesis while maintaining the bioactivity of HATG (HATG-Alg bioink, Fig. 2A) in parallel to the expansion of hAUR in the presence of GFs. Due to the calcium triggered crosslinking (CTEC) mechanism of HATG (*38*), crosslinking of Alg and HATG can be initiated simultaneously by the diffusion of calcium ions (Fig. 1B).

**Fig. 2.**
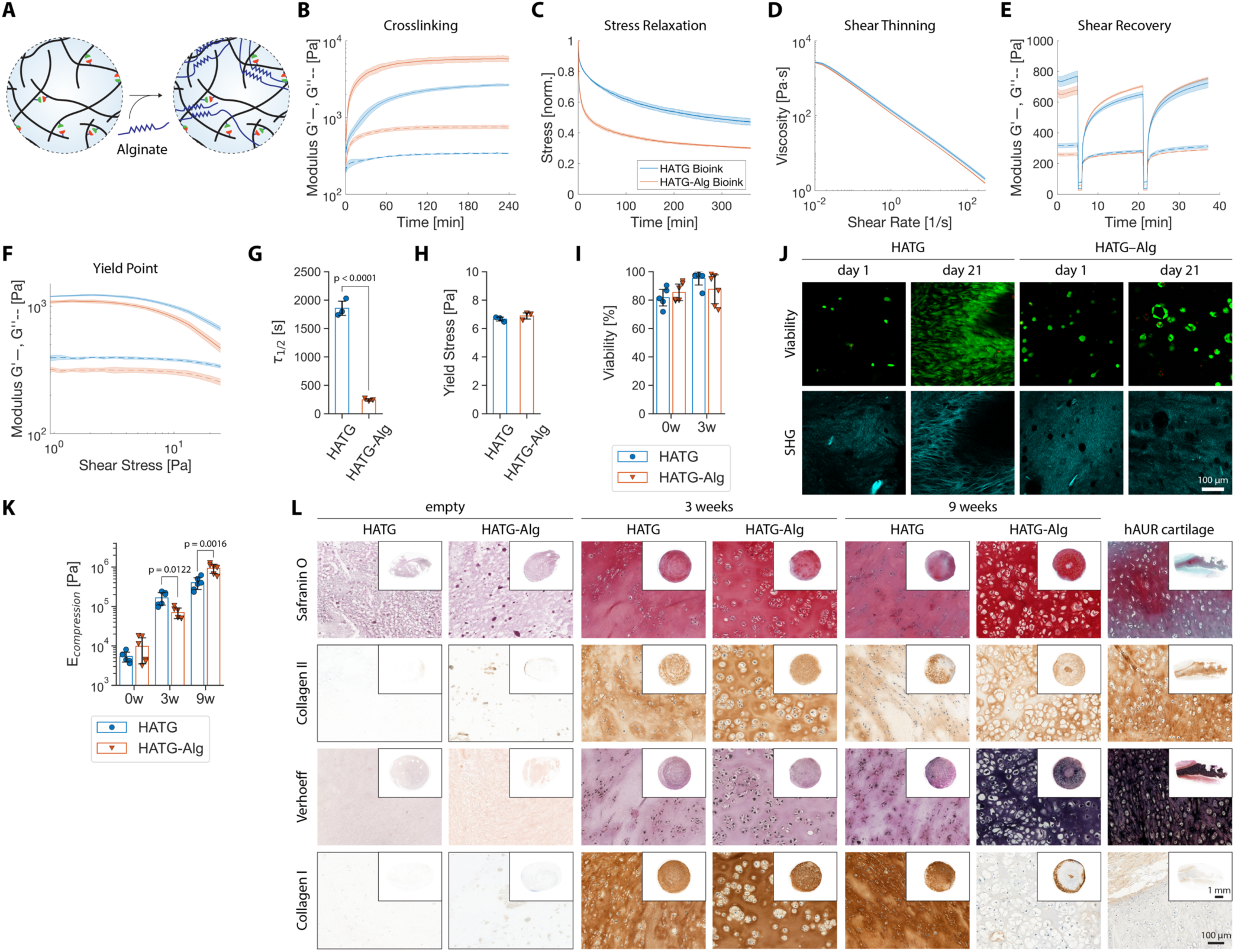
Adaptation of the HATG bioink through the addition of alginate results in the formation of elastic cartilage. (**A**) Illustration of the HATG bioink (left) and HATG-Alg bioink (right). Cell expansion was performed in the absence of additional growth factors and 10 million cells per mL were used. (**B-F**) Rheology of crosslinking (**B**), stress relaxation (**C**), shear thinning (**D**), shear recovery (**E**) and stress sweep (**F**) of the HATG and HATG-Alg bioink. (**G**) Relaxation halftime defined by the time the bioink requires to relaxed to half of the initial minus final stress from stress relaxation tests (**C**). (**H**) Yield stress calculated from stress sweep tests (**E**). (**I**) Viability of hAUR encapsulated in the HATG and HATG-Alg bioink at a density of 10 million cells per mL after 1 day and 3 weeks. (**J**) Representative viability images (live: green, dead: red) of hAUR in the HATG and HATG-Alg bioink after 1 day and 3 weeks as well as second harmonic generation (SHG) images. Scale bar: 100 µm. (**K**) Compressive modulus of grafts after 1 day and 3 and 9 weeks. (**L**) Histology of grafts after 3 and 9 weeks compared to empty bioink gels and human auricular (hAUR) cartilage as control. Scale bar: close up: 100 μm, inserts: 1 mm. n_d_ = 2, n_s_ = 6.

The HATG-Alg bioink resulted in stiffer gels reaching 5.8 ± 0.5 kPa in storage modulus (G’) compared to 2.7 ± 0.1 kPa of the HATG bioink (Fig. 2B), despite reducing the HATG concentration from 1.5% to 0.5% to reduce stiffness. While the HATG bioink was softer than the HATG-Alg bioink, previous studies showed that a higher HATG concentration (> 2%) detrimentally impacts chondrogenesis (*53*). We therefore opted not to match the stiffness of the HATG bioink to the stiffness of the HATG-Alg bioink. Both bioinks displayed similar shear thinning, shear recovery and yielding behavior (Fig. 2D-F). Stress-relaxation tests confirmed that the addition of Alg decreases the stress relaxation half-time significantly (HATG: 1857 ± 125 s, HATG-Alg: 243 ± 21 s, Fig. 2C,G).

Upon encapsulation of human auricular (hAUR) chondrocytes in either bioink, cells displayed distinctly different morphologies with cells being spread and elongated in the HATG bioink, whereas they maintained a round, chondrogenic morphology in the HATG-Alg bioink (Fig. 2I,J). Upon maturation for 9 weeks, HATG-Alg grafts reached compressive moduli of 0.9 ± 0.2 MPa (Fig. 2K), approaching values of native human auricular cartilage (1.2 ± 0.1 MPa). HATG grafts, in contrast, reached only half of the stiffness of HATG-Alg grafts (0.4 ± 0.1 MPa). The addition of Alg further led to the deposition of elastin in almost all samples (5 out of 6), compared to only 2 out of 6 samples in HATG grafts without alginate. Collagen I, a marker for fibrocartilage, was absent in all HATG-Alg grafts, except for a thin collagen I ring around the edges of the grafts, potentially related to a self-organizing perichondrium (*54*). In contrast, collagen I was present throughout the entire graft in half of the HATG grafts (Fig. 2L). Glycosaminoglycans (GAGs) and collagen II displayed stainings similar to native human auricular cartilage (Fig. 2L).

The addition of Alg therefore substantially improved the maturation of our grafts towards elastic rather than fibrocartilage, reaching a composition and stiffness approaching native human auricular cartilage. The round morphology of chondrocytes in the HATG-Alg bioink supports the role of alginate in improving chondrogenesis through mechanical confinement (*52*). However, differences in bioink stiffness could impact chondrogenic behavior as well, requiring further investigation into the origin of the fibro- to elastic cartilage transition by reducing HATG and Alg concentrations in the HATG-Alg bioink to match the stiffness of the HATG bioink.

### Increasing cell density results in grafts maturing into juvenile elastic cartilage

We then combined the expansion of chondrocytes in the presence of GFs with the alginate modification of the bioink to further improve graft maturation. Despite their individually positive effects, combining the addition of GFs during expansion and the modification with alginate negatively impacted the onset of elastic cartilage formation observed in the HATG-Alg bioink (Fig. 2) and resulted in 2-fold softer grafts (Fig. S5). In addition, inhomogeneities in ECM deposition could be observed, possibly related to the lower cell viability in constructs with cells expanded in the presence of GFs.

To understand why grafts matured inhomogeneously and were substantially softer, we analyzed the cell density in areas of impaired tissue maturation and compared it to the cell density in areas of good tissue maturation (Fig. 1E). Areas of impaired tissue maturation had a 3 to 4-fold lower cell density compared to areas of good tissue maturation, underlining previous results which showed that a reduction in cell density is detrimental to tissue maturation (Fig. S6). We therefore increased the cell density from 10 to 30 million cells per mL to account for impaired tissue maturation in areas of low cell density (Fig. 3A).

**Fig. 3.**
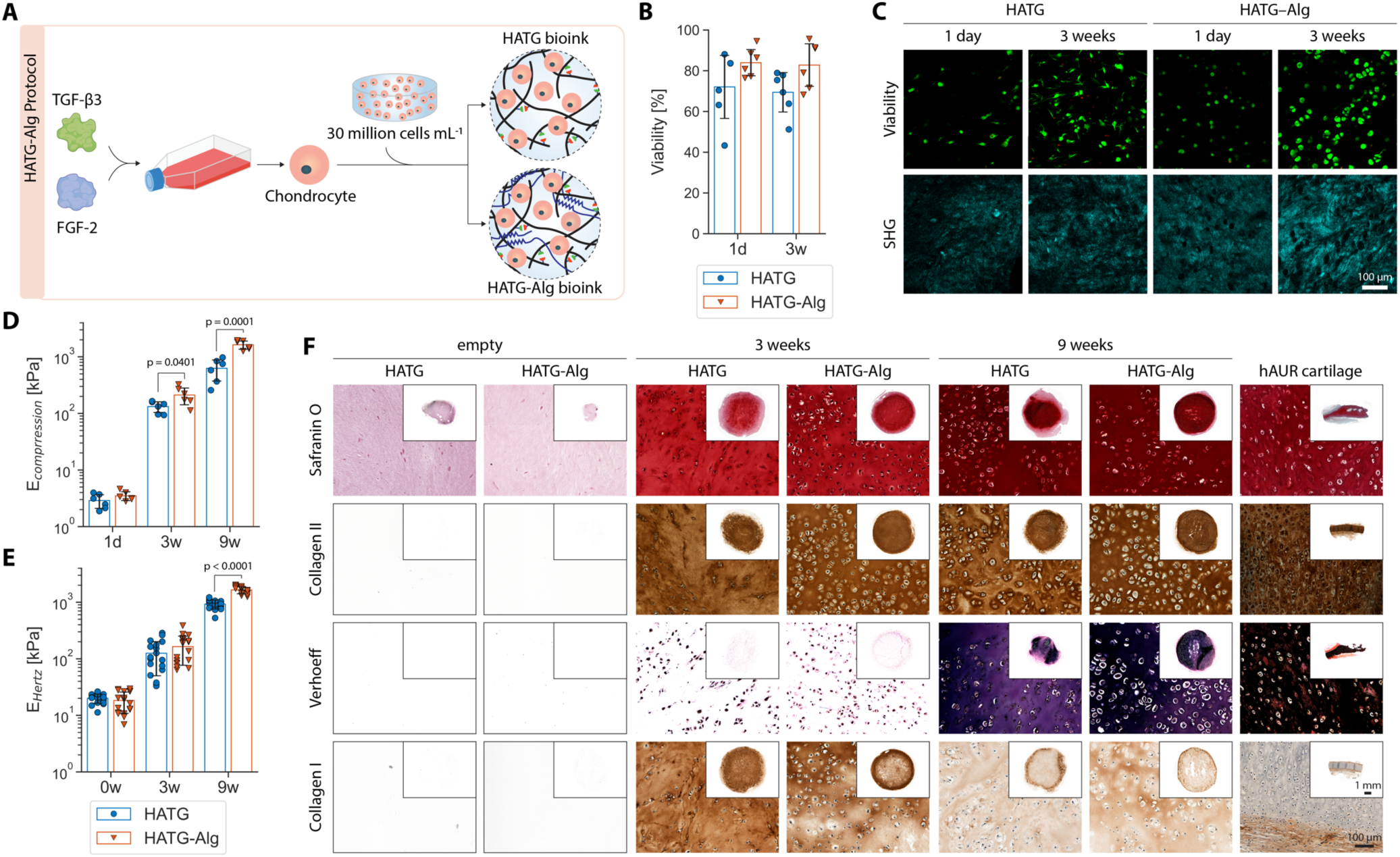
Elastic cartilage formation in bioprinted grafts. (**A**) hAUR chondrocytes were expanded in the presence of FGF-2 and TGF-β3 and combined with either the HATG or HATG-Alg bioink at a cell density of 30 million cells per mL. (**B**) Cell viability in the HATG and HATG-Alg bioink 1 day and 3 weeks after printing. (**C**) Representative images of cell viability (green = viable, red = dead) and second harmonic generation for either bioink after 1 day and 3 weeks. Scale bar: 100 µm. (**D**) Compressive and (**E**) Hertz modulus of grafts after 1 day, 3 and 9 weeks. (F) Histology of grafts after 3 and 9 weeks compared to empty bioink samples and human auricular (hAUR) cartilage as control. Brightness adjusted Verhoeff staining – original Verhoeff staining depicted in Fig. S7. Scale bar: close up: 100 μm, inserts: 1 mm. n_d_ = 2, n_s_ = 3.

The increase in cell density substantially improved graft maturation, resulting in grafts maturing homogeneously throughout. ECM deposition after 9 weeks of culture closely resembled that of native human auricular cartilage in histological sections (Fig. 3F). Functionally, HATG-Alg grafts reached compression and indentation moduli of 1.5 ± 0.3 MPa (Fig. 3D,E), matching the mechanical properties of native human auricular cartilage (1.2 ± 0.1 MPa) and exceeding the moduli obtained after GF expansion (0.6 ± 0.1 MPa, Fig. S4C) and alginate supplementation (0.9 ± 0.2 MPa, Fig. 2K) alone. HATG grafts in contrast reached only half the stiffness of HATG-Alg (compression: 0.6 ± 0.3 MPa, indentation: 0.9 ± 0.2 MPa).

### Effect of cumulative improvements on gene expression demonstrates robust shift from fibrocartilage to mature cartilage

To evaluate the cumulative impact of the improved protocol, we performed RNA-seq on grafts generated using the original (HATG (*38*), fibrocartilage) and improved (HATG-Alg, elastic cartilage) protocol at 3 and 9 weeks (Fig. 4A, Fig. S8-S11). The cumulative changes resulted in a robust upregulation of genes associated with cartilage matrix formation and chondrogenic identity, including key ECM components (COL2A1, COL9A1-3, COL11A2, MATN3, MATN4, CNMD) and cartilage-regulatory factors (GREM1, FGFR3, CSGALNACT1, PAPSS2). Correspondingly, genes linked to ossification (DMP1, BGLAP), angiogenesis (EREG, VCAM1, F3), neurogenesis (NES, GABRB2, SLC6A4), and inflammation (RSAD2, TNFSF10, CSF1) were significantly downregulated (Fig. 4B,C).

**Fig. 4.**
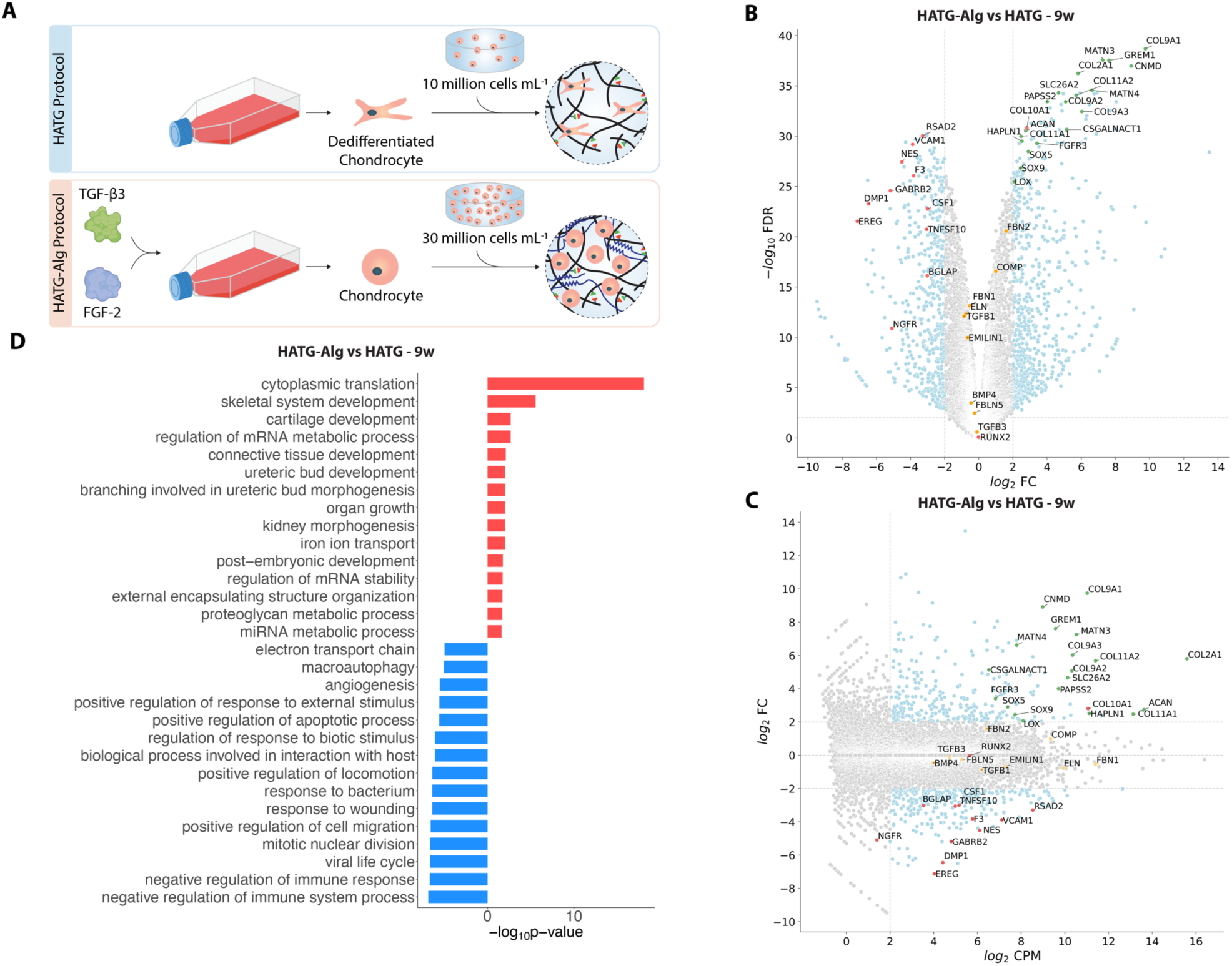
Transcriptional evaluation of hAUR in grafts prepared following the HATG protocol compared to the HATG-Alg protocol. (**A**) HATG protocol: hAUR were expanded in the absence of additional growth factors and combined with the HATG bioink at a density of 10 million cells per mL. HATG-Alg protocol: hAUR were expanded in the presence of 10 ng mL^-1^ FGF-2 and TGF-β3 and combined with the HATG-Alg bioink at a density of 30 million cells per mL. (**B-C**) Volcano and MA plots comparing samples of the HATG-Alg against samples of the HATG protocol after 9 weeks of culture. Green: upregulated genes positively influencing cartilage formation, red: genes negatively influencing cartilage formation and/or involved in other pathways, orange: further genes related to elastic cartilage formation. (**D**) Gene ontology (GO) analysis of samples of the HATG-Alg against the HATG protocol after 9 weeks of culture (red: upregulated in HATG-Alg, blue: downregulated in HATG-Alg). n_d_ = 1, n_s_ = 3.

To further explore the functional implications of these transcriptional shifts, we performed a gene ontology (GO) enrichment analysis (Fig. 4D, Fig. S11). In agreement with our analysis of differentially expressed genes, upregulated genes in HATG-Alg grafts were enriched for processes related to skeletal system development, cartilage development, proteoglycan metabolism and ECM organization. Downregulated genes were enriched for pathways related to immune responses, angiogenesis, neurogenesis and wound healing.

Taken together, the gene expression profiles and GO enrichment data support the transition away from fibrocartilage towards a more mature cartilage identity. Although markers for elastic cartilage such as ELN and FBN1 were highly expressed (logCPM > 10) in both conditions, the histological presence of elastin in HATG-Alg grafts and absence in HATG grafts suggests additional regulatory mechanisms may contribute to elastin protein deposition, potentially influenced by the presence of genes upregulated in HATG-Alg grafts. These findings demonstrate that the cumulative improvements in cell expansion, bioink design, and culture conditions promote the formation of auricular cartilage approximating native elastic cartilage, even in the absence of transcriptional changes in key elastic fiber genes.

### Lysyl oxidase improves collagen but not elastin fibril network crosslinking

Lysyl oxidase (LOX), a copper dependent amine oxidase, catalyzes the formation of inter- and intra-molecular crosslinks in collagen and elastin fibrils. These crosslinks are essential for fiber maturation, contributing to the maturation of cartilage (*55, 56*) and resistance to enzymatic degradation (*57*). With the lack of copper in many culture media, LOX activity is impaired. Previous studies have shown the substantial impact LOX and copper supplementation has on hyaline cartilage graft stiffness (*55, 58*). We therefore supplemented our culture media with exogenous lysyl oxidase-like protein 2 (LOXL2) and/or copper to evaluate its impact on graft stiffness and crosslinks (Fig. S12). Notably copper alone was sufficient to significantly increase collagen crosslinks (pyridinoline (PYD) and deoxypyridinoline (DPD), Fig. S12S) and improve graft indentation modulus (Fig. S12E). However, the compressive modulus remained unchanged (Fig. S12F) and elastin crosslinks (desmosine (DES) and isodesmosine (isoDES)) remained below the detection limit (Fig. S12T). In addition, a trend towards reduced tensile and bending moduli was observed (Fig. S12H,I), despite comparable histological appearance (Fig. S12U). While changes in elastin crosslinks under the addition of copper and/or LOXL2 might occur but remain undetected due to the lower detection limit, this limit was substantially lower than the elastin crosslinks observed in human auricular cartilage. The lack of elastin crosslinks therefore suggests an immature elastic fiber network. Combined with the naturally low concentration of collagen II in elastic cartilage, this may explain the reduced tensile and bending stiffness as collagen II alone may be insufficient to create a tissue-spanning fiber network. Nevertheless, given the role of crosslinked protein networks in protecting against enzymatic degradation (*59–61*), we continued to supplement our culture media with copper for our *in vivo* experiments.

### Bioprinted elastic cartilage grafts demonstrate *in vivo* stability

To assess *in vivo* performance, we bioprinted cylindrical grafts using the HATG-Alg protocol (Fig. 5A), matured them for 9 weeks *in vitro*, and implanted them subcutaneously in immunocompromised rats for an additional 6 weeks (Fig. 5B). Similar to our previous experiments, hAUR remained viable post-printing (Fig. 5C-D) and grafts matured into elastic cartilage-like tissue over the time course of 9 weeks (Fig. 5F-Q). Both instantaneous and equilibrium moduli matched those of human auricular cartilage (grafts: E_instantaneous_ = 1.1 ± 0.03 MPa, E_equilibrium_ = 68 ± 10 kPa, human auricular cartilage: E_instantaneous_ = 1.0 ± 0.1 MPa, E_equilibrium_ = 81 ± 11 kPa, Fig. 5F-G). Grafts reached a dry weight of 11 ± 2.5% (human auricular cartilage: 27 ± 1.4%, Fig. 5H) with GAGs (Fig. 5I), collagen II (Fig. 5K) and LOX-mediated collagen crosslinks (Fig. 5M) matching those of human auricular cartilage while collagen I remained below the detection limit (Fig. 5L). Histological stainings confirmed these observations (Fig. 5Q).

**Fig. 5.**
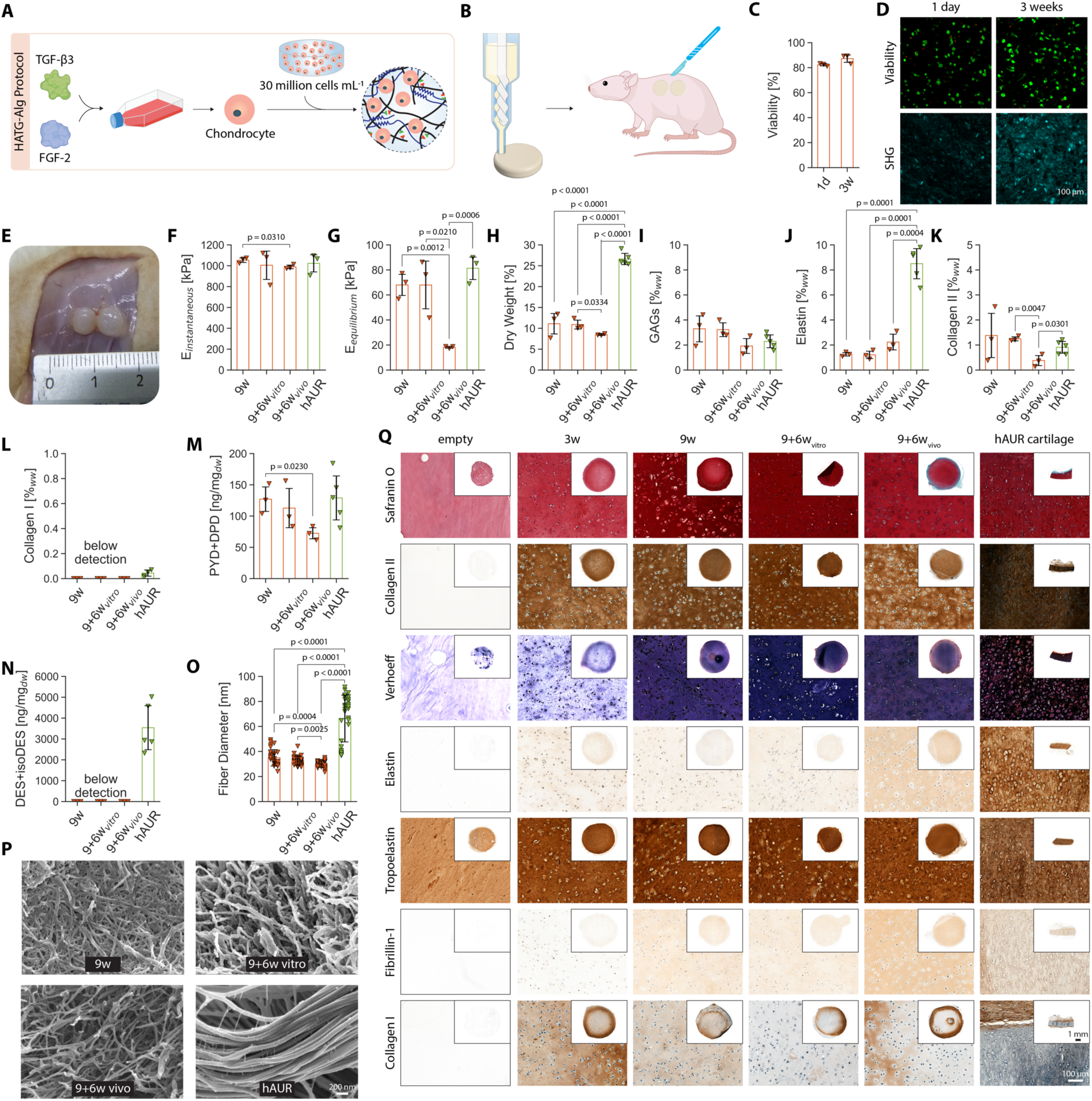
Tissue engineered elastic cartilage-mimetic grafts are stable *in vivo*. (**A**) Illustration of the HATG-Alg protocol used to promote the maturation of grafts into elastic cartilage. (**B**) Grafts were bioprinted using a progressive cavity pump-based extruder (Fig. S13) and implanted subcutaneously after 9 weeks of maturation in immunocompromised rats for 6 weeks. (**C**) Cell viability 1 day and 3 weeks after bioprinting. (**D**) Representative images of cell viability and second harmonic generation. Scale bar: 100 µm. (**E**) Image of grafts after 6 weeks *in vivo*. (**F-G**) Instantaneous and equilibrium modulus. (**H-L**) Evaluation of graft ECM composition: dry weight and GAGs, elastin, collagen II and collagen I (detection limit: 0.027 ± 0.01%_dw_) as percentage wet weight (%_ww_ – percentage dry weight depicted in Fig. S14). (**M-N**) LOX mediated crosslinks in collagen (pyridinoline and deoxypyridinoline, PYD+DPD) and elastin (desmosine and isodesmosine, DES+isoDES, detection limit: DES: 117 ng mg_dw_^-1^, isoDES: 160 ng mg_dw_^-1^). (**O-P**) SEM images of tissue engineered elastic cartilage and evaluation of their fiber diameter. Scale bar: 200 nm. (**Q**) Histological and immunohistological evaluation of tissue engineered elastic cartilage compared to human auricular cartilage and empty HATG-Alg samples. Brightness adjusted Verhoeff staining – original Verhoeff staining depicted in Fig. S14. 9w: 9 weeks *in vitro* culture, 9+6w vitro: 9+6 weeks *in vitro* culture, 9+6w vivo: 9 weeks *in vitro* culture followed by 6 weeks *in vivo*. Scale bar: zoom in: 100 μm, full view: 1 mm. n_d_ = 1, n_s_ = 3.

Grafts further showed the deposition of elastin (Fig. 5Q), with an elastin content of 1.3 ± 0.1%_ww_ (Fig. 5J), which was significantly lower than that of human auricular cartilage (8.5 ± 1.0%_ww_). We therefore examined the maturation state of the elastin network. Elastogenesis is a complex process involving the coordinated expression of multiple proteins for the formation of mature elastic fibers (*56*). The two main components of elastic fibers are fibrillin, which forms microfibrils, and tropoelastin, which aligns along the microfibrils to form mature elastic fibers. Immunohistological stainings for elastin, tropoelastin and fibrillin-1 revealed a pronounced deficiency in fibrillin-1 as main cause for the lack of mature elastic fibers (Fig. 5Q), confirming our observations in Verhoeff-stained sections lacking large elastic fiber bundles (Fig. 5Q). These findings were further supported by the lack of LOX-mediated elastin crosslinks (Fig. 5N). In addition, the smaller fiber diameter and visually shorter fiber length observed in SEM images indicates an immature protein network, although no distinction between elastin and collagen II can be made(Fig. 5O-P).

Grafts however remained structurally stable *in vivo* (Fig. 5E) maintaining both their instantaneous modulus (Fig. 5F) and dry weight (Fig. 5H). Histologically, a stronger staining for fibrillin-1 as well as elastin was observed after explantation (Fig. 5Q) and elastin content increased (Fig. 5J), reaching a composition similar to human auricular cartilage in percent dry weight (Fig. S14). Although total elastin per wet weight remained lower, these findings suggest continued elastin network maturation *in vivo*. The drop in equilibrium modulus and smaller fiber diameter indicate that this process is incomplete.

### Bioreactor platform enables uniform maturation of auricular grafts

We next assessed whether bioprinted auricular grafts would mature into elastic cartilage (Fig. S15A-E). After 9 weeks, ears developed an opaque appearance and became flexible and elastic. However, compressive (center: 92 ± 112 kPa, edge: 556 ± 231 kPa) and indentation (center: 40 ± 51 kPa, edge: 961 ± 674 kPa) moduli differed significantly between the center and the edge of the auricle. Consistent with the lower mechanical properties, histological analysis showed elastic cartilage-like tissue at the periphery, while central regions resembled fibrocartilage. As a result, grafts lacked sufficient mechanical integrity and collapsed *in vivo* (Fig. S15F-H).

To improve structural integrity, we therefore aimed to promote a homogeneous development of these grafts. A key factor in tissue development is nutrient accessibility, which strongly influences tissue maturation (*62, 63*). Studies have further shown that TGF-β accumulates on the surface of grafts while central parts remain devoid of it (*63, 64*). To address this, we designed a bioreactor platform that suspends auricular grafts in an elevated position, enabling nutrient diffusion from all directions (Fig. 6A). This setup resulted in a uniform maturation, with significantly improved mechanical properties across the construct (center: 740 ± 75 kPa, edge: 867 ± 150 kPa, Fig. 6B-C). In addition, no differences in histological stainings were observed between center and edge (Fig. 6P).

**Fig. 6.**
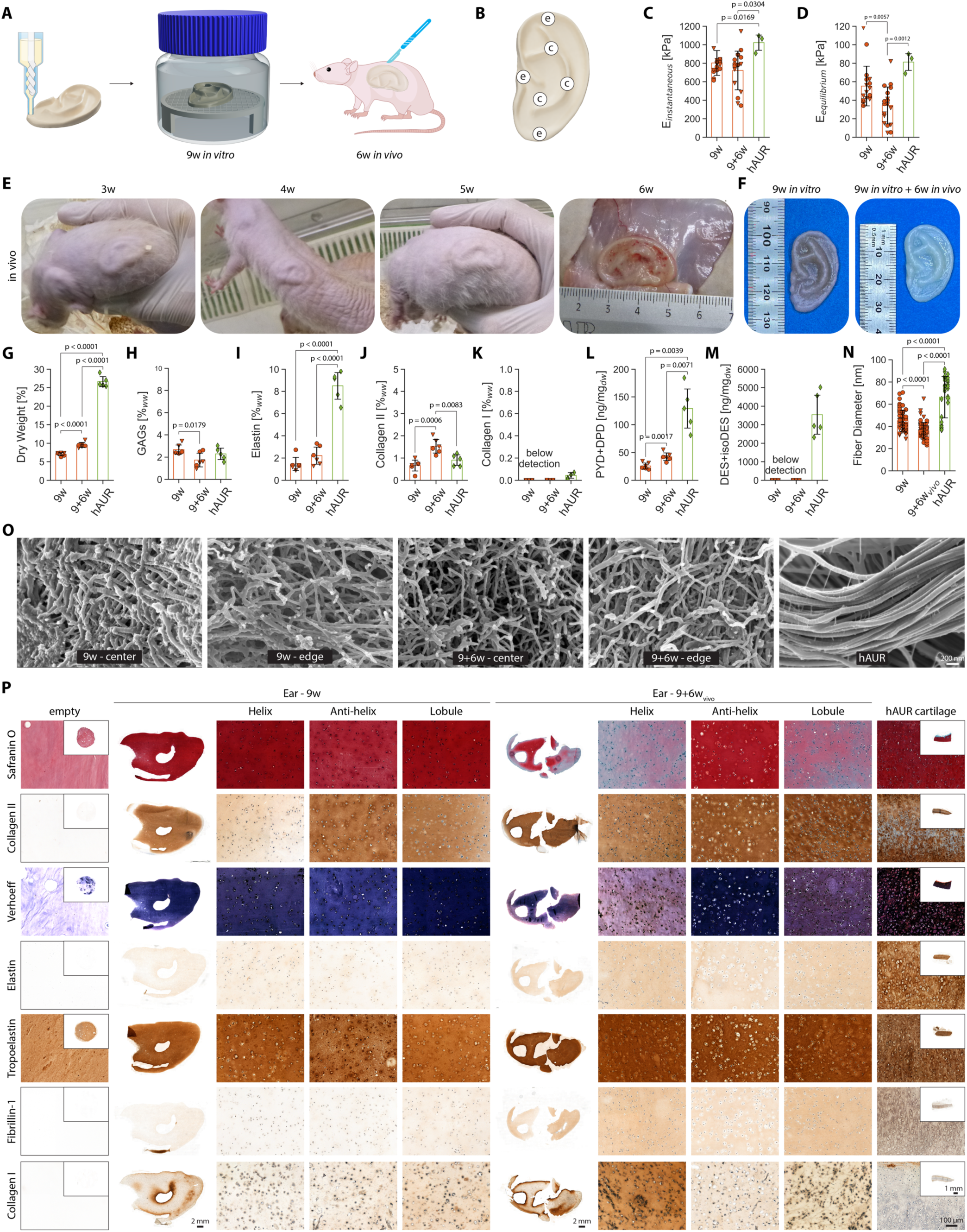
*In vivo* stability of tissue engineered elastic cartilage-mimetic auricular grafts. (**A**) Auricular grafts were bioprinted using a progressive cavity pump-based extruder, cultured on a custom designed bioreactor platform to allow nutrient perfusion from all sides of the graft for 9 weeks, and implanted subcutaneously in immunocompromised rats for 6 weeks. (**B**) Positions of the auricular graft where compression tests were carried out (e: edge, c: center). (**C-D**) Instantaneous and equilibrium modulus. (**E**) Images of auricular grafts implanted subcutaneously in immunocompromised rats. (**F**) Images of auricular grafts after 9 weeks of *in vitro* maturation (9w *in vitro*) and an additional 6 weeks *in vivo* (9w *in vitro* + 6w *in vivo*). (**G-K**) Evaluation of graft ECM components: dry weight and GAGs, elastin, collagen II and collagen I (detection limit: 0.027 ± 0.01%_dw_) as percentage wet weight (%_ww_ – percentage dry weight depicted in Fig. S17). (**L-M**) LOX mediated crosslinks in collagen (pyridinoline and deoxypyridinoline, PYD+DPD) and elastin (desmosine and isodesmosine, DES+isoDES, detection limit: DES: 117 ng mg_dw_^-1^, isoDES: 160 ng mg_dw_^-1^). (**N-O**) SEM images of tissue engineered elastic cartilage and evaluation of their fiber diameter. Scale bar: 200 nm. (**P**) Histological and immunohistological evaluation of tissue engineered elastic cartilage compared to human auricular cartilage and empty HATG-Alg samples. Brightness adjusted Verhoeff staining – original Verhoeff staining depicted in Fig. S17. 9w: 9-week culture, 9+6w: 9-week culture plus 6 weeks *in vivo*. Scale bar: zoom in: 100 μm, full view: 1 mm. n_d_ = 1, n_s_ = 3.

### Elastic cartilage-mimetic auricular grafts exhibit lasting elasticity and structural integrity *in vivo*

Once a homogeneous maturation was achieved, we implanted auricular grafts subcutaneously in immunocompromised rats to simulate placement beneath the temporal scalp as performed in microtia reconstruction. The grafts retained their shape and contour beneath the skin (Fig. 6A,E) and showed no deformations after 6 weeks (Fig. S16C-F). The grafts further displayed elastic behavior regaining their original shape after being bent or twisted (Fig. S16G, Movie S1). Upon explantation, grafts were encapsulated by a thin, vascularized layer of fibrous tissue, while the grafts themselves remained avascular.

Compositionally, auricular grafts approached the properties of native human elastic cartilage, with comparable levels of GAGs (Fig. 6H), collagen II (Fig. 6J) and absence of collagen I (Fig. 6K). Dry weight (Fig. 6G), elastin (Fig. 6I) as well as collagen and elastin crosslinks (Fig. 6L-M) however remained below levels of human auricular cartilage, indicating a more immature protein network. Histologically, grafts exhibited a uniform deposition of GAGs, collagen II, and elastin (Fig. 6P). Although mature elastic fibers were not observed histologically and in SEM images (Fig. 6N-P), immunostaining for tropoelastin, fibrillin-1 and elastin demonstrated a progressive *in vivo* maturation, indicated by the increased staining intensity for elastin and fribrillin-1 (Fig. 6P, Fig. S17F-H). Importantly neither calcification nor vascular invasion was observed (Fig. S17I-J).

While longer implantation periods are known to permit elastic fiber maturation (*32–35, 65*), the peripheral degradation of GAGs, elastin, and collagen II observed after 6 weeks (Fig. 6P) highlights the need to further strengthen the elastic fiber network. Such a fully mature, crosslinked elastin architecture will further enhance resistance to enzymatic degradation (*59–61*) and ensure the long-term structural stability of these grafts. In summary, these findings demonstrate that tissue engineered auricular grafts can achieve structural stability and lasting elasticity *in vivo*. Together with the transition towards mature elastic fibers, these results define a clear path towards clinically viable, patient-specific ear reconstruction.

## Discussion

The ability to create auricular grafts from a small (∼5 mg) biopsy, eliminating the need for costal cartilage harvest, has the potential to provide microtia patients with a durable, real, and natural ear. With the successful maturation of auricular grafts with near-native elastic cartilage properties before implantation, we have achieved a critical milestone in replicating the auricle for the treatment of microtia. The maturation of these grafts before implantation eliminates the reliance on internal (*34–36, 65*) or external (*32, 33*) support scaffolds to sustain physiological forces and the inhomogeneous maturation in an uncontrolled *in vivo* environment (*34–37*).

Previous efforts in auricular tissue engineering have largely pursued two strategies: seeding chondrocytes onto synthetic polymer frameworks, typically based on polyglycolic acid (PGA) and polylactic acid (PLA) meshes (*34–36*) or encapsulating them within hydrogels (*31, 37*). Mesh frameworks offer immediate mechanical support, and mesh reinforcements can be incorporated through polycaprolactone (PCL) scaffolds, but cell seeding is inefficient and matrix deposition thus inhomogeneous (*34, 35*). Moreover, while mechanically tunable, these synthetic frameworks do not reproduce the natural elasticity of the auricle. In contrast, hydrogel-based approaches facilitate homogeneous cell distribution and extracellular matrix formation, yet their intrinsic softness leads to graft deformation *in vivo* (*31, 37*). Without appropriate cartilage maturation, approaches to stabilize these grafts using external scaffolds are required to improve shape retention (*32, 33*). However, these approaches do not address graft maturation and rely on the *in vivo* environment to act as bioreactor. This leads to uncontrolled and inhomogeneous tissue maturation with areas showing the formation of elastic cartilage, while others remain devoid of it (*31–33*).

The formation of elastic fibers has been reported after prolonged *in vivo* maturation (*32–35, 65*), but elastic fibers were typically sparsely distributed and occurred only localized, with the bulk of these grafts consisting of other tissue. The inability to create mature cartilage prior to implantation, and the uncontrolled *in vivo* environment resulting in only localized graft maturation in contrast to the tissue-spanning elastic fiber network observed in auricular cartilage therefore results in graft deformation if not addressed by internal (*34–36, 65*) or external (*32, 33*) support scaffolds.

Recent work combined a nanoparticle-based Runx1 plasmid delivery to dedifferentiated auricular chondrocytes with double-network hydrogels, yielding cartilage-like tissue with EvG-positive elastic fibers and improved mechanics after 1-3 months *in vivo* (*66*). In contrast, our gene-free multifactorial approach employing the maintenance of the chondrogenic potential during expansion, the modulation of the stress relaxation behavior, and the increase in cell density and graft maturation, matures auricular grafts *ex vivo* to near-native elastic cartilage with uniform elastin, collagen II and GAG distribution, and absence of collagen I, before implantation, thereby reducing reliance on *in vivo* remodeling. While the approaches we employed have been studied individually, mainly in articular cartilage, their combination in an integrated approach for the maturation towards near-native tissue properties presents a substantial progress to move the field beyond temporary shape retention towards the generation of functional elastic cartilage.

To support long-term performance and enable clinical translation, we identified elastogenesis, as remaining step towards fully functional elastic cartilage. Elastogenesis is a tightly regulated, multifactorial process involving the expression, assembly and alignment of tropoelastin and fibrillins into mature elastic fibers (*67*). While the assembly process has been studied extensively, knowledge gaps regarding the specific genetic programs and ECM interactions that regulate this process, particularly in elastic cartilage, remain. For example, BMP-4 has been shown to enhance elastin and fribillin-1 production in synergy with TGF-β1 in cardiac relevant cell types (*68*). Post-translationally, fibrillins organize into microfibrils, which serve as scaffolds for coacervated tropoelastin assembly (*67*). This process can be impaired by GAGs through steric hindrance or inhibition by heparan sulfate (*69–71*), suggesting that the temporal modulation of the GAG-matrix e.g. via enzymatic degradation may promote microfibril assembly, similar to its effect on the collagen II network formation in hyaline cartilage (*58, 72*). Concurrently, mechanosensitive pathways play a critical role in stimulating elastic fiber synthesis and maturation (*73–75*). A deeper understanding of the spatiotemporal regulation of these processes during auricular development may guide strategies to engineer fully functional elastic cartilage. Notably, the maturation of elastic fibers *in vivo* underscores the feasibility to generate fully functional elastic cartilage in tissue engineered auricular grafts.

Regarding the translational potential of this study, all raw components are available in GMP-grade, including sNAG nanofibrils, alginate, hyaluronan, peptides and growth factors. The synthesis of HATG, however, will require the development of a GMP-compliant production process. Beyond the raw materials, translation will also require a reproducible scale-up of graft fabrication, validation of bioprinting under GMP conditions, and the integration of quality control measures for cell expansion, matrix evaluation and mechanical testing for product release. Particularly the combination of cells and biomaterials is likely to be classified as an advanced therapy medicinal product (ATMP) subject to stringent regulatory requirements.

The primary human auricular chondrocytes used in this study represent a clinically relevant cell source for the treatment of microtia. Once established, our protocols reproducibly supported the maturation of grafts into elastic cartilage in six independent experiments using cells from five human donors (age: 8.2 ± 4.4 years, range: 5–15 years; sex: 3 female, 2 male), underscoring the robustness of the approach. In unilateral microtia cases, chondrocytes from the unaffected ear could serve as an autologous source, provided genetic testing excludes mutations in key developmental regulators such as HOXA2 or BMP pathway genes (*76–79*). Broader genetic heterogeneity has been reported in syndromic microtia, with variants identified in TCOF1 (Treacher Collins syndrome), FGF3 (LAMM syndrome), or DHODH (Miller syndrome) (*1, 80*). Although many of these associations arise in syndromic contexts or small cohorts, they underscore the genetic heterogeneity of microtia and the importance of targeted genetic testing to guide safe and effective use of patient-specific cells. In bilateral microtia, alternative sources such as microtic, nasal, or costal chondrocytes may be required, potentially complemented by induced pluripotent stem cells and genetic engineering.

While promising, several limitations must be addressed before these grafts can be applied in patients. First, the ear grafts in this study were implanted for six weeks, far shorter than the duration required in humans. Before advancing to extended *in vivo* studies, peripheral graft degradation underscores the need for a mature elastic network to ensure long-term stability. These long-term studies will further be required to exclude risks such as calcification or abnormal tissue remodeling. Second, we used an immunocompromised rat model, which facilitates evaluation of human cells but does not capture the immune dynamics of autologous transplantation. Testing in autologous immunocompetent models such as rabbit (*40*) or sheep (*65, 81*) will be essential, potentially requiring strategies to manage inflammatory responses during early healing. Third, our use of auricular chondrocytes from otoplasty procedures does not fully replicate the intended therapeutic scenario, where donor cells may derive from healthy or microtic ears with possible genetic defects. Finally, surgical translation must be considered: whereas our grafts were implanted subcutaneously, clinical reconstruction involves the elevation of the auricle in a second stage, often requiring additional cartilage wedges and skin grafting. Skin contraction and scarring remain risks to the initial shape fidelity until fully healed. Achieving faithful elastic cartilage formation is therefore a prerequisite not only for aesthetic restoration but for long-term graft durability and mechanical compatibility with the native auricle. Given the shared compositional and mechanical features among cartilaginous tissues, this strategy may also be adaptable to reconstructive applications involving tracheal, nasal and articular cartilage, where mechanical resilience and structural fidelity are equally essential.

In conclusion, our auricular grafts offer a path to personalized therapy of microtia without the complications associated with autologous costal cartilage reconstruction, polyethylene implants, or osseointegrated epitheses. The grafts presented in this study represent the closest approximation of native elastic cartilage achieved *ex vivo* to date. We anticipate that this work will lay the foundation for the development of fully mature, clinically viable elastic cartilage, enabling a durable, long-term therapy for children affected by microtia.

## Materials and Methods

### Cell Source

Primary human auricular chondrocytes were isolated from cartilage remnants of otoplasty surgery. 11 donors were used in total in this work (patient age: 8.4 ± 3.9 years, age range: 5–17 years, sex: 6 female, 5 male). 6 donors were used to establish the protocols to mature grafts into elastic cartilage (patient age: 8.7 ± 4.1 years, age range: 6–17 years, sex: 3 female, 3 male), while 5 donors were used to mature grafts into elastic cartilage (patient age: 8.2 ± 4.4 years, age range: 5–15 years, sex: 3 female, 2 male). Informed consent was obtained from the patient or their legal guardian under the ethical license BASEC-Nr.2017–02101. Approval of all experimental protocols was obtained from the Ethics Committee of the Canton Zurich (Kantonale Ethikkommission Zürich). Due to ethical and institutional regulations, these human-derived materials are not available for distribution to third parties.

### Human Auricular Chondrocyte Isolation and Culture

Two protocols were used for the isolation of human auricular chondrocytes, described below with reference to the respective data and figures presented in this study.

For the experiments depicted in Fig. S1 and Fig. S6, the following protocol was used. Auricular cartilage remnants were washed twice in PBS containing 50 μg mL^-1^ gentamicin (Gibco) before pre-digestion with pronase solution (DMEM 31966, 0.5% v/v pronase (pronase from streptomyces, Roche), 10% v/v fetal bovine serum (FBS, Gibco) and 1x Antibiotic-Antimycotic (Gibco)) at 37°C for 90 min. After digestion, samples were washed again in PBS containing 50 μg mL^-1^ gentamicin, freed from the tissue surrounding them and the cartilage minced into 1-2 mm pieces. Pieces were then digested in collagenase solution (DMEM, 10% v/v FBS, 1x Antibiotic-Antimycotic, 0.1% collagenase from clostridium histolyticum (C6885, Merck, CDU: collagen digesting units)) at 37°C overnight. Cells were collected by passing the resulting solution through a 40 μm cell strainer. The obtained solution was centrifuged at 250 rcf for 10 min and the cell pellet collected. Cells were expanded by plating them at a density of 10’000 cells cm^-2^ in DMEM containing 20% v/v FBS, 50 μg mL^-1^ L-ascorbate-2-phosphate (TCI) and 1x Antibiotic-Antimycotic to passage 1. Expansion was carried out in standard tissue culture flasks (TPP, T150) at 37°C, 5% CO_2_ and 95% humidity.

For all remaining experiments the following isolation protocol was used. Auricular cartilage remnants were washed twice in PBS containing 50 μg mL^-1^ gentamicin (Gibco) before pre-digestion with pronase solution (DMEM 31966, 40 U mL^-1^ pronase (pronase from streptomyces, Roche), 2% v/v fetal bovine serum (FBS, Gibco) and 1x Antibiotic-Antimycotic) at 37°C for 90 min. After digestion, samples were washed again in PBS containing 50 μg mL^-1^ gentamicin, freed from the tissue surrounding them and the cartilage minced into 1-2 mm pieces. Pieces were then digested in collagenase solution (DMEM, 2% v/v FBS, 1x Antibiotic-Antimycotic, 1000 CDU mL^-1^ collagenase from clostridium histolyticum (C6885, Merck, CDU: collagen digesting units)) at 37°C overnight. Cells were collected by passing the resulting solution through a 40 μm cell strainer. The obtained solution was centrifuged at 250 rcf for 10 min and the cell pellet collected. Cells were expanded by plating them at a density of 10’000 cells cm^-2^ in DMEM containing 10% v/v FBS, 50 μg mL^-1^ L-ascorbate-2-phosphate (TCI), 1x Antibiotic-Antimycotic and 5 ng mL^-1^ FGF-2 (Recombinant Human FGF-basic, PeproTech) to passage 1. Expansion was carried out in standard tissue culture flasks (TPP, T150) at 37°C, 5% CO_2_ and 95% humidity.

From passage 1 onwards, cells were expanded by seeding them at a density of 3’000 cells cm^-2^ to passage 3. Cells were expanded in the presence of DMEM containing 10% v/v FBS and 10 µg mL^-1^ gentamicin (basal medium). In addition, different expansion protocols were tested: FGF-2: 10 ng mL^-1^ FGF-2 were added to the basal medium. FGF-2+BMP-2: 10 ng mL^-1^ FGF-2 and 10 ng ml^-1^ BMP-2 (Recombinant Human/Murine/Rat BMP-2, PeproTech) were added to the basal medium. FGF-2+BMP-9: 10 ng mL^-1^ FGF-2 and 10 ng ml^-1^ BMP-9 (Recombinant Human GDF-2, PeproTech) were added to the basal medium. FGF-2+TGF-β3: 10 ng mL^-1^ FGF-2 and 10 ng ml^-1^ TGF-β3 (Recombinant Human TGF-β3, PeproTech) were added to the basal medium. At confluency, cells were trypsinized, combined with basal medium and centrifuged at 250 rcf for 4 min. The cell pellet was collected and either reseeded or washed with TRIS buffered glucose (TBG; 50 × 10^−3^ M TRIS, 200 × 10^−3^ M d-glucose, pH 7.6) to prevent any CaCl_2_ from being mixed into the bioink. Cells were mixed with the respective bioinks at a concentration of 10 and 30 million cells mL^-1^.

### Peptide Synthesis

Transglutaminase substrate peptides (glutamine donor peptide: H-NQEQVSPL-ERCG-NH2, lysine donor peptide: Ac-FKGG-ERCG-NH2) were synthesized on a peptide synthesizer (Prelude X, Gyros Protein Technologies AB). Peptides were synthesized on Rink Amide MBHA resin (RAM-100-HL, Gyros Protein Technologies AB) at a 0.69 mmol g^-1^ loading capacity and yield of 800 μmol per reaction vessel. The resin was allowed to swell in dimethylformamide (DMF) for 1 h followed by deprotection in 20 mL of 20% pieridine in DMF at 50°C for 1 min twice and three washing steps in DMF. The first amino acid was coupled at 2 eq. in 2 eq. hexafluorophosphate benzotriazole tetramethyl uronium (HBTU) and 4 eq. N-methyl morpholine (NMM) at 50°C for 5 min, and the coupling step repeated three times with DMF washing in between. The following amino acids were coupled by deprotection as mentioned before and coupling at 2 eq. in 2 eq. HBTU and 4 eq. NMM at 50°C for 5 min, and the coupling step repeated twice with DMF washing in between. At the end of the synthesis, the last amino acid was deprotected and the FKGG-ERCG was acetylated using 20% acetic anhydride in DMF and washed four times in DMF. Finally, peptides were washed with dichloromethane (DCM) six times and dried for 1h under nitrogen. Peptides were cleaved in 20 mL cleavage solution containing 4% water, 4% triisopropylsilane (TIPS) and 2% EDDT in trifluoroacetic acid (TFA) for 2 h at room temperature and afterwards precipitated in cold diethyl ether.

Synthesized peptides were purified on a preparative HPLC system (Agilent) using an Agilent Prep LC column (C18, 10 µm particle size, 100 Å pore diameter, 250 mm length, 50 mm diameter, AG410910-502, Agilent). Gradient runs (0-3 min: 10% solvent B, 3-13 min: gradient 10% to 50% solvent B, 13-15 min: gradient 50% to 90% solvent B, 15-16 min: 90% solvent B - solvent A: 0.1% TFA in water, solvent B: 0.1% TFA in acetonitrile) were performed at a flow rate of 80 mL min^-1^. Retention peaks from previously successful syntheses were used as reference. Successful peptide synthesis was further confirmed by mass spectrometry.

### HATG Synthesis

Hyaluronan transglutaminase (HATG) was synthesized as previously reported (*38, 82*). Briefly, hyaluronan (HA, Lifecore Biomedical) was thiolated with 3,3’-dithiobis(propanoic dihydrazide) in a first step. In a second step, vinyl sulfonated hyaluronan was created by the addition of divinyl sulfone. Lastly, peptide modification was carried out by letting either the glutamine donor peptide (NQEQVSPL-ERCG) or the lysine donor peptide (FKGG-ERCG) react to vinyl sulfonated hyaluronan. The final product was filtered through a 0.2 μm filter and finally lyophilized. A degree of substitution of ∼10-13% over the disaccharide repeating units was measured by NMR. Successful synthesis was further confirmed through rheological crosslinking measurements comparing crosslinking dynamics and plateau storage moduli to previously successful batches.

### Bioink Preparation

The HATG bioink was prepared as reported previously (*38*). sNAG nanofibrils (Marine Polymer Technologies, US) were dissolved in TRIS buffered glucose (TBG, 200 × 10^-3^ M d-glucose (Merck), 50 × 10^-3^ M Tris (Merck), pH 7.6) at a concentration of 4.82%. Fibril dissociation was ensured by sonication (Branson 2510EMTH). In a first step, the sNAG solution was combined with a 1.15% w/w HA in TBG solution. Afterwards, HA and HATG were stepwise added to avoid agglomerations of nanofibrils. Once mixed with the cell volume and enzymes, the final concentrations were 1.5% HATG, 1.5% w/w HA and 2% sNAG. In a similar fashion, the HATG-Alg bioink was prepared to reach final concentrations of 0.5% HATG, 0.25% Alg, 1.5% HA and 2% sNAG. sNAG nanofibrils were obtained from Marine Polymer Technologies Inc. Before printing either bioink was combined with hAUR at their respective concentration and FXIII and thrombin to reach a final concentration of 20 U mL^-1^ FXIII (Fibrogammin, CSL Behring) and 1 U mL^-1^ Thrombin (TISSEEL, Baxter).

### Gel Casting

Either bioink was casted into UV-sterilized PDMS molds (SYLGARD 184, Corning) of 4 mm diameter and 1 mm height. Crosslinking was initiated by adding 2 mL of 100 × 10^-3^M CaCl_2_ for 1 h. Samples were cultured in chondrogenic medium consisting of DMEM 31966, 50 µg mL^−1^ L-ascorbate-2-phosphate, 40 µg mL^−1^ L-proline, 10 µg mL^−1^ gentamycin, 1% ITS+ (ITS+ Premix Universal Culture Supplement, Corning) and various growth factors at a concentration of 10 ng mL^-1^. If no information on the exact growth factor added is stated, the growth factor added was TGF-β3. Culture was performed in 12 well-plates with 2 mL of media changed every 3-4 days.

### Bioprinting

Bioprinting was performed on a Biofactory bioprinter (regenHU, Switzerland) enclosed in a laminar flow hood. STL models were sliced in Slic3r (https://slic3r.org) and post-processed in a custom written Matlab postprocessor (MATLAB 2019a, MathWorks). Bioinks were laden into 3 or 10 cc printing cartridges and briefly centrifuged. A pneumatic extrusion system was initially used for bioprinting. Printing was performed with a 410 μm conical nozzle, 10 mm s^-1^ moving speed and at a pressure of approximately 45 kPa. Later, the pneumatic extrusion system was exchanged to a progressive cavity pump-based system (Puredyne Kit B, ViscoTec Pumpen- u. Dosiertechnik GmbH) interfaced with the bioprinter through an Arduino controller coupled to a TMC2209 stepper motor driver. Constructs were printed onto glass plates which were subsequently transferred into a 100 × 10^-3^ M CaCl_2_ solution for crosslinking. Crosslinking was carried out for 1 h in case of cylinders (6 mm diameter, 1 mm height) or for 2 h in case of auricular constructs. After crosslinking, constructs were transferred into culture flasks and cultured in chondrogenic medium consisting of DMEM 31966, 50 µg mL^−1^ L- ascorbate-2-phosphate, 40 µg mL^−1^ L-proline, 10 µg mL^−1^ gentamycin, 1% ITS and 10 ng mL^-1^ TGF-β3, changed every 3-4 days. In the case of copper and/or LOX supplementation, 2.5 µg mL^-1^ copper sulfate (Merck) and 0.15 µg mL^-1^ lysyl oxidase-like protein 2 (SignalChem) were supplemented.

### Viability

Samples were briefly washed in DMEM 31966 and subsequently stained in DMEM 31966 supplemented with 1 × 10^-6^ M Calcein AM (Invitrogen) and 1 × 10^-6^ M propidium iodide (PI, Fluka) for 1 h. Afterwards samples were washed again and imaged on a multiphoton microscope (Leica SP8 MP) equipped with a 25x water immersion objective. Calcein AM and PI excitation was performed at 900 nm (Mai Tai XF, Spectra-Physics). Calcein AM emission was collected from 500 to 570 nm and PI emission was collected from 695 to 740 nm. Simultaneously, second harmonic generation (SHG) was collected from 442 to 458 nm. SHG was confirmed by moving the slider of the detector out of the described range and a sharp signal loss observed. Z-Stacks were imaged from the samples surface up to 100 μm into the sample at an interval of 1 μm between stacks. Viability was analyzed by counting viable and dead cells in images at the surface and at a distance of 25, 50, 75 and 100 μm from the surface and by dividing the number of viable cells by the number of total cells.

### Rheology

Rheology was performed on an MCR301 and MCR302e rheometer (AntonPaar) equipped with a Peltier element (P-PTD 200, AntonPaar) and thermal hood (H-PTD 200, AntonPaar). Tests were carried out with a 20 mm parallel plate geometry (PP20, AntonPaar) at 25°C. A wet tissue was placed around the sample in the thermal hood to prevent dehydration of the sample. A poly-L-lysine (PLL) coating was applied to the measurement geometry to prevent slippage between sample and geometry. A 10 μg mL^-1^ PLL solution was placed between the two plates of the rheometer and the temperature raised to 37°C. After 30 min, the PLL solution was removed and the samples loaded.

Crosslinking was tested in oscillatory tests at 1% shear strain, 1 Hz and a gap of 200 µm. Data points were collected every 20 s and tests performed for 4 h. After 1 min, crosslinking was initiated by placing 0.5 mL of 100 × 10^-3^ M CaCl_2_ solution around the sample to allow CaCl_2_ to diffuse in and initiate crosslinking.

Shear thinning tests were carried out in rotational tests by logarithmically increasing the shear rate from 0.01 s^-1^ to 300 s^-1^ while data point collection was decreased logarithmically from 300 s per datapoint to 1 s per datapoint. Tests were performed at a gap of 200 µm.

Shear recovery was tested in 5 intervals in oscillatory mode at a gap of 600 µm and 1 Hz. In a first interval, the material was tested at low shear (1% shear strain) for 300 s, followed by a second interval at high shear (500% shear strain) for 60 s. Afterwards a brief 5s resting interval followed and the recovery measured in a third interval at 1% shear strain for 900 s. The high-shear, 5 s rest and recovery interval were then repeated once.

To calculate the yield point shear stress amplitude sweeps from 0.5 to 25 Pa were carried out at a gap of 600 µm at 1 Hz. Yield stress analysis was performed in Rheoplus (v3.62, AntonPaar) using the Yield Stress I analysis.

Stress relaxation tests were carried out by first crosslinking the samples at a gap of 500 µm for 4 h between the two plates as in the crosslinking tests and then applying a shear strain of 15%. Samples were then allowed to relax for 6 h. The shear stress was normalized to its initial value and the relaxation half time (τ_½_) calculated by calculating the time, the material needed to relax to half of the initial stress minus final stress value.

### Compression Testing

Unconfined compression, tensile and bending tests were performed on a TA.XTplus Texture Analyzer (Stable Micro Systems) equipped with a 500 g load cell.

Compressive modulus (E_compression_): Samples were placed between the two compression plates and a pre-load applied to ensure proper contact between the sample and the plates. Samples were allowed to relax for 5 min and testing was performed at 0.01 mm s^-1^ up to 15% strain. Loading and unloading curves were recorded and the compressive modulus calculated from the slope of the first 3% of the loading curve.

Stress-relaxation (E_instantaneous_ and E_equilibrium_): Samples were placed between the two compression plates and a pre-load (0.005 N for empty gels, 0.05 N for 3-week-old samples and 0.25 N for 9-week-old samples) applied to ensure proper contact between the sample and the plates. Samples were allowed to relax for 10 min and 4 consecutive stress-relaxation intervales applied. During each interval, the strain was increased by 5% at a strain rate of 5% s^-1^ to reach a final strain of 20% after the 4^th^ interval. During the first two intervals, samples were allowed relax for 1800 s and after the last two intervales for 2700 s. To calculate the instantaneous modulus (E_instantaneous_), a linear fit of the loading curve between 15% and 20% strain was applied. The equilibrium modulus (E_equilibrium_) was calculated by fitting a line through the stress and strain values after relaxation of each interval.

Bending modulus (E_bending_): Sample dimensions were recorded using a stereomicroscope (Leica Wild Heerbrugg). Samples were then placed on a 3-point bending setup (Three Point Bend Rig – small, Stable Micro Systems) with a gap of 10 mm and radius of 6 mm. Samples were bent at 0.02 mm s^-1^. Bending stress and strain were calculated according to ISO 178:2019 and the modulus determined by linear fitting the first 10% strain of the bending stress-strain curve.

Tensile modulus (E_tensile_): Sample dimensions were recorded using a stereomicroscope (Leica Wild Heerbrugg). Samples were then clamped in a tensile setup (Miniature Tensile Grips, Stable Micro Systems) using sandpaper to prevent slipping during the test. The sample length was determined at the start of each test and samples extended at 0.1 mm s^-1^. The tensile modulus was calculated by linear fitting the first 10% of the tensile stress-strain curve.

### Bioindentation Testing

Indentation tests were performed on an UNHT3 Bio Bioindenter (AntonPaar) equipped with a spherical ruby indenter (Ø 1 mm). Samples were fixed on petri dishes with super glue. Once the glue hardened samples were submerged in 0.9% NaCl and tested. Samples were first approached with the indenter to identify their surface upon which they were indented for either 25, 40 or 60 μm depending on the samples stiffness as to avoid overloading the loadcell of the bioindenter within 5 s. Stress relaxation was recorder for 5 min after which the samples were unloaded. 3 measurements were performed at each measurement point per sample. Hertz modulus (E_Hertz_) was calculated with the Indentation software (Indentation 8.0.15, AntonPaar).

### Subcutaneous Rat Implantation

Subcutaneous immunocompromised rat implantation was performed in accordance with the ethical license ZH147/2019 approved by the Veterinary Office of the Canton Zürich (Veterinäramt, Gesundheitsdirektion, Kanton Zürich, Switzerland) and experiments conducted following the Swiss regulatory framework (ethical principles and guidelines for experiments on animals, Swiss Academy of Sciences and Swiss Academy of Medical Sciences, 2005) and the ETH Zurich policy on experimental animal research.

2 months old male immunocompromised rats (Rj:ATHYM-Foxn1^rnu/rnu^) were obtained from Javier Labs. 30 min prior surgery analgesia was administered (Meloxicam, Metacam, 2 mg kg^−1^, subcutaneous). Anesthesia was initiated by 4–5% isoflurane and eye cream applied to prevent desiccation of the cornea. Anesthesia was maintained at 1.5–3%. An incision along the dorsal midline was created and a pocket created by blunt dissection. Auricular grafts were placed subcutaneously and the incision closed with surgical staples. A wound sealing spray was applied to seal the incision. One-week post-surgery staples were removed. After 6 weeks rats were euthanized by CO_2_ asphyxiation, samples collected and further processed.

### Histology and Immunohistochemistry

Constructs were fixed in 4% paraformaldehyde for 4 h and then dehydrated in graded ethanol solutions. Samples were paraffin-embedded and 10 μm sections cut on a microtome. Samples were deparaffinized and rehydrated before stainings. Brightfield imaging was performed on an automated slide scanner (Panoramic 250, 3D Histech).

#### Safranin O

Slides were stained in Weigert’s iron hematoxylin solution for 5 min, washed in deionized water and differentiated in 1% acid alcohol (1% of 37% HCl in 70% ethanol) for 2 s and washed again in deionized water. Following slides were stained in 0.02% Fast Green for 1 min, destained in 1% acetic acid for 30 s, and stained in 1% Safranin O for 30min. Slides were then dehydrated to xylene and mounted.

#### Verhoeff’s stain

Slides were stained according to the manufacturers protocol (HT25, Merck). Briefly, slides were stained in working elastic stain solution for 10 min, rinsed in deionized water and differentiated in working ferric chloride solution for 30 s. Afterwards slides were rinsed in deionized water again, dehydrated to xylene and mounted. To better visualize elastic fibers in stained sections, brightness of the images was adjusted in ImageJ, ensuring that each image of the same staining, i.e. samples, and positive and negative controls, were adjusted the same way. Unadjusted images are displayed in the supplementary figures.

#### Immunohistochemistry

Antigen retrieval was carried out by treating slides with hyaluronidase (8000 U mL^−1^) at 37°C for 30 min. Slides were washed in PBS and blocked with 5% BSA in PBS at room temperature. Afterwards slides were treated with the primary antibody in 1% BSA in PBS at 4°C overnight (primary antibodies: Table 1). Slides were then washed and treated with 0.3% H_2_O_2_ for 15 min. Afterwards slides were washed again and the secondary antibody in 1% BSA in PBS applied for 1 h (secondary antibodies: Table 1). Slides were washed again and developed with the DAB substrate kit (ab64238, Abcam) according to the manufacturer’s specifications for 5 min. Slides were then washed, stained in Weigert’s iron hematoxylin solution for 3 min, destained in 1% acid alcohol for 2 s, blued in 0.1% Na_2_CO_3_ for 1 min, dehydrated to xylene and mounted. Experiments were grouped and stained together with the same positive (human auricular cartilage) and negative (empty gels) controls.

**Table 1.** Immunohistochemistry antibodies and dilutions.

| Target | Primary antibody | Secondary antibody |
| --- | --- | --- |
| Collagen I | Anti-Collagen I (EPR7785) (0.875 mg mL<sup>-1</sup>, 1:1500, ab138492, Abcam) | Goat Anti-Rabbit IgG H&L (HRP) (2 mg mL<sup>-1</sup>, 1:1000, ab6721, Abcam) |
| Collagen II | Anti-Collagen II (II-II6B3) (75 µg mL<sup>-1</sup>, 1:20, DSHB Hybridoma Product II-II6B3, deposited by Linsenmayer, T.F.)(83) | Goat Anti-Mouse IgG H&L (HRP) (2 mg mL<sup>-1</sup>, 1:1000, ab6789, Abcam) |
| Elastin | Anti-Elastin (ELN/1981) (0.2 mg mL<sup>-1</sup>, 1:200, NBP2-75767, Novus Biologicals) | Goat Anti-Mouse IgG H&L (HRP) (2 mg mL<sup>-1</sup>, 1:1000, ab6789, Abcam) |
| Tropoelastin | Anti-Tropoelastin (1:200, PR398, Elastin Product Company) | Goat Anti-Rabbit IgG H&L (HRP) (2 mg mL<sup>-1</sup>, 1:1000, ab6721, Abcam) |
| Fibrillin-1 | Anti-Fibrillin 1 (FBN1/6933) (0.2 mg mL<sup>-1</sup>, 1:200, NBP3-20532, Novus Biologicals) | Goat Anti-Mouse IgG H&L (HRP) (2 mg mL<sup>-1</sup>, 1:1000, ab6789, Abcam) |

#### Semi-Quantitative Evaluation

Semi-quantitative evaluation of immunohistological stainings for elastin, tropoelastin and fibrillin-1 was carried out as reported previously (*38*). Color deconvolution and measurement of the mean gray value of the resulting images were carried out in ImageJ (Gabriel Landini, https://imagej.net/Colour_Deconvolution, v3.0.2). After deconvolution, mean gray values were inverted and the average value of empty samples subtracted. All images were analyzed the same way. Vectors for the deconvolution are presented in Table 2.

**Table 2.** Color vectors used for the semi-quantitative evaluation of immunohistological stainings for elastin, tropoelastin and fibrillin-1.

| Staining | Vector | Red | Green | Blue |
| --- | --- | --- | --- | --- |
| Elastin | 1 | 0.23440556 | 0.48759046 | 0.841017 |
|  | 2 | 0.5190854 | 0.56108177 | 0.6447772 |
|  | 3 | 0.31490505 | 0.47751275 | 0.82025385 |
| Tropoelastin | 1 | 0.17157604 | 0.3970757 | 0.90160555 |
|  | 2 | 0.41717747 | 0.5358375 | 0.734058 |
|  | 3 | 0.28273103 | 0.40888208 | 0.8676858 |
| Fibrillin-1 | 1 | 0.14226002 | 0.45548728 | 0.87880224 |
|  | 2 | 0.54620063 | 0.5818733 | 0.6025681 |
|  | 3 | 0.22490622 | 0.2735681 | 0.9351886 |

### RT-qPCR

RNA isolation was performed using NucleoZOL according to the manufacturer’s specifications (Machery-Nagel) after 2D expansion in 6-well plates. RNA concentration was determined using a nanodrop reader (Thermo Scientific) and 1 µg used for reverse transcription quantitative polymerase chain reaction (RT-qPCR). The GoTaq qPCR Master Mix (Promega) was used for PCR amplification and reactions were run on a QuantStudio 3 96-well 0.1 ml Block Real-Time PCR System (ThermoFisher). The primers used in this study are listed in Table 3. Data analysis was carried out using the 2^-ΔΔCt^ method and normalized against the housekeeping gene GAPDH.

**Table 3.**
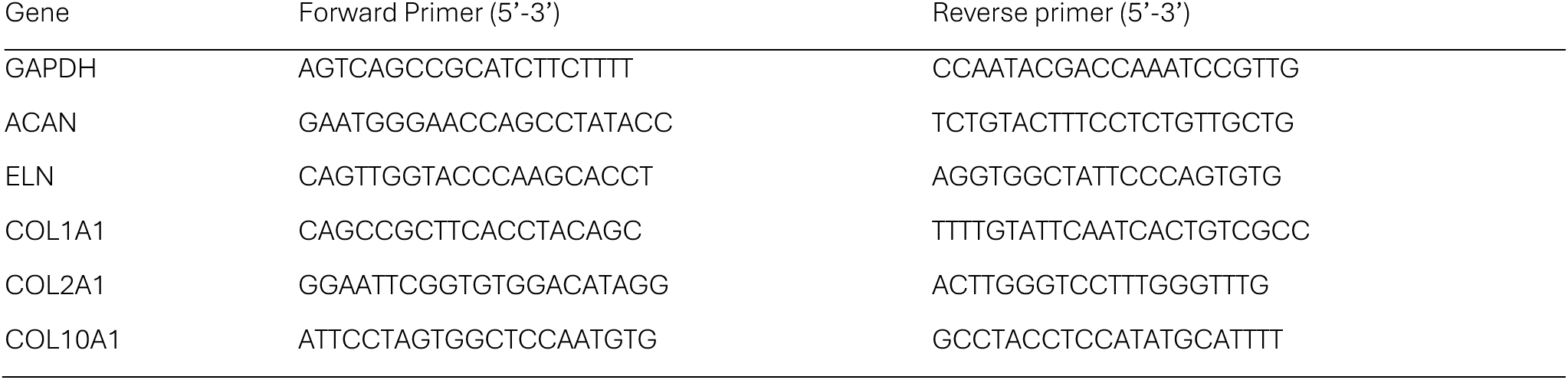
q-PCR primer list.

### ECM quantification

#### Dry weight

To determine the dry weight of grafts, grafts were weighted, placed in Eppendorf tubes, lyophilized for 48 h and weighted again. The dry weight percentage was calculated as weight after lyophilization (dry weight) divided by the weight before lyophilization (wet weight). ECM components are represented as percentage dry weight (%_dw_) as well as percentage wet weight (%_ww_).

#### Glycosaminoglycans

GAGs were determined using the Blyscan (Biocolor Ltd) assay kit. 1-2 mg of lyophilized sample was taken and digested in papain extraction reagent (270 µg mL^-1^ papain, 8 mg mL^-1^ sodium acetate, 4 mg mL^-1^ EDTA, 0.8 mg mL^-1^ cysteine in 200 × 10^-3^ M sodium phosphate buffer, pH 6.4) overnight at 65°C. To enhance digestion, samples were mechanically dissociated using a positive displacement pipette after 3 h in the papain extraction reagent. Samples were aspirated into a 50 µL pipette tip, the tip pressed against the Eppendorf tube walls, and the sample extruded, creating shearing. After digestions, samples were allowed to cool to room temperature, centrifuged at 13000 rcf for 10 min and the supernatant collected. 5-100 µL of sample, dependent on the GAG concentration, were taken, sample volume adjusted to 100 µL using deionized water, combined with 1 mL Blyscan dye reagent and incubated for 30 min. Afterwards, samples were centrifuged at 13000 rcf, the tube’s liquid drained, 1 mL dissociation reagent added to release the bound dye and incubated for 10 min. Samples were centrifuged again at 13000 rcf for 10 min and 100 µL of the dye extract from each sample pipetted into a 96-well flat bottom well plate. Samples were read on a microplate reader (Synergy H1, BioTek) at 656 nm. GAG concentration for each sample was calculated using the reference standards and blank.

#### Elastin

Elastin was determined using the Fastin (Biocolor Ltd) assay kit. 1-2 mg of lyophilized sample was taken and extracted in 750 µL of 250 × 10^-3^ M oxalic acid at 100°C for 60 min. The entire extraction process was repeated 5 times to extract all elastin as determined with human auricular cartilage samples in advance. After each extraction step, samples were cooled to room temperature, centrifuged at 13000 rcf for 10 min and the supernatant collected. After the first extraction, samples were mechanically dissociated as described in the glycosaminoglycan assay. The exact volume of the extraction was determined using a pipette and 50-250 µL taken for the assay. An equal volume of elastin precipitating reagent was added to the sample and incubated for 15 min. Samples were centrifuged at 13000 rcf for 10 min, the tube’s liquid drained, 1 mL of dye reagent added and incubated on a shaker for 90 min. Afterwards, samples were centrifuged at 13000 rcf for 10 min, the entire tube’s liquid carefully drained, 250 µL of dye dissociation reagent added and incubated for 10 min. 100 µL of the dye extract from each sample were pipetted into a 96-well flat bottom well plate and samples were read on a microplate reader (Synergy H1, BioTek) at 513 nm. Elastin concentration for each sample was calculated using the reference standards and blank.

#### Collagen I and II

Collagen I and II were determined using the Type I and Type II Collagen Detection ELISA Kits from Chondrex (Chondrex Inc). 1-2 mg of lyophilized sample was taken and pretreated in 500 µL distilled water at 4°C overnight. The samples were spun down at 13000 rcf for 3 min and the supernatant discarded. The samples were then placed in 500 µL 3 M guanidine dissolved in 50 × 10^-3^ M Tris-HCL, pH 7.5, at 4°C overnight to remove GAGs and enhance pepsin digestion. Samples were centrifuged at 13000 rcf for 3 min and then washed in distilled water followed by an overnight wash in 500 µL of 50 × 10^-3^ M acetic acid at 4°C. Pepsin digestion was carried out in 500 µL pepsin solution (0.1 mg mL^-1^ pepsin in 50 × 10^-3^ M acetic acid) on a shaker at 4°C for 48 h, repeated 5 times. After the second extraction, samples were mechanically dissociated as described in the glycosaminoglycan assay each time the pepsin solution was changed. After each extraction step sample solution was collected in cold 200 µL buffered normal goat serum (100 × 10^-3^ M Tris-base, 150 × 10^-3^ M NaCl, pH 7.5). In a final step, elastin was digested in 500 µL pancreatic elastase (0.1 mg mL^-1^ in 100 × 10^-3^ M Tris-base, 150 × 10^-3^ M NaCl, 5 × 10^-3^ M CaCl_2_, pH 7.8) at 4°C for 24h. Samples were centrifuged at 13000 rcf and the supernatant added to the collection tube containing normal goat serum. After digestion, the exact volume of the extraction was determined using a pipette and 1/50 of the final volume of 1 M Tris-base was added to the supernatant. All steps were performed on ice to prevent collagen from gelling.

After digestion, ELISA plates were prepared by incubation with 100 µL of capture antibody solution at 4°C overnight. 100 µL of standards and diluted samples (dilution determined during assay setup) were then added and incubated for 2 h. Afterwards, 100 µL of detection antibody was added and incubated for 2 h, followed by the addition of 100 µL streptavidin peroxidase solution for 1 h. 100 µL of OPD solution were then added, incubated for 30 min and stopped by the addition of 50 µL stop solution. Samples were read on a microplate reader (Synergy H1, BioTek) at 490 nm and collagen I and II concentration for each sample was calculated using the reference standards and blank. Three plate washes were performed between each step.

#### Collagen crosslinks

Pyridinoline (PYD) and deoxypyridinoline (DPD) crosslinks were determined by hydrolyzing approximately 1 mg of sample (dry weight) in 6 M HCl at 110°C for 48 h. Samples were then dried in a vacuum concentrator for 24 h and diluted in water to reach a final concentration of 10 mg mL^-1^. HPLC runs were performed on a Jasco HPLC system using a TSKgel ODS-80Tm HPLC reverse phase column (C18, 5 µm particle size, 100 Å pore diameter, 150 mm length, 4.6 mm diameter, 808148, Merck). Isocratic runs of 0-40 min solvent A (0.13% heptafluorobutyric acid (HFBA), 24% methanol), 40-50 min solvent B (0.1% HFBA, 75% acetonitrile), 50-60 min solvent A were performed at a flow rate of 0.8 ml min^-1^ (pump: PU-2080 Plus, Jasco). Samples were diluted 5 times in 0.5% HFBA, 10% acetonitrile in water to reach a sample concentration of 2 mg mL^-1^. 25 µL of sample, corresponding to 50 µg sample dry weight, were injected for each run (autosampler: AS-2055 Plus, Jasco). Fluorescence was recorded at 400 nm (excitation: 295 nm, detector: FP-2020 Plus, Jasco). A PYD/DPD standard (8004, Quidel Corporation) was used to quantify PYD/DPD crosslinks in grafts(*55*).

#### Elastin crosslinks

Desmosine (DES) and isodesmosine (isoDES) crosslinks were determined by hydrolyzing approximately 1 mg of sample (dry weight) in 6 M HCl at 110°C for 48 h. Samples were then dried in a vacuum concentrator for 24 h and diluted in water to reach a final concentration of 10 mg mL^-1^. HPLC runs were performed on an Agilent HPLC system with UV detector (1260 DAD WR, Agilent) using a PolySULFOETHYL A ion exchange column (C18, 5 µm particle size, 300 Å pore diameter, 35 mm length, 2.1 mm diameter, 3.52SE0503, PolyLC Inc). Gradient runs (0-6 min: 100% solvent A, 6-7 min: gradient 0% to 52.5% solvent B, 6-7 min: 52.5% solvent B, 7-8 min: gradient 52.5% to 0% solvent B, 8-10 min: 100% solvent A – solvent A: 0.1% TFA in water, solvent B: 1 M NaCl, 0.1% TFA in water) at a flow rate of 0.36 mL min^-1^ were performed (pump: 1260 Quat Pump, Agilent). Samples were diluted 1:2 in water to reach a final sample concentration of 5 mg mL^-1^ and 20 µL of sample injected, corresponding to 100 µg sample dry weight (autosampler: 1260 Vialsampler, Agilent). UV absorption at 240 nm and 268 nm was recorded to determine DES and isoDES crosslinks. DES (c_des_) and isoDES (c_iso_) concentration was determined using the following formula:

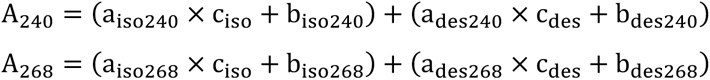

where A is the area under the peak and a and b are molar absorption coefficients of DES and isoDES at 240 nm and 268 nm respectively. A DES and isoDES standard (DES: DD866, isoDES: MS264, Elastin Product Company) was used to quantify DES and isoDES crosslinks in grafts(*84*).

### RNA-sequencing

#### RNA extraction

Grafts were snap frozen in liquid nitrogen and subjected to cryopulverization. The powder was collected in 1.4 mL TRIzol Reagent (Invitrogen, 15596026), and RNA was isolated according to the manufacturer’s specifications. RNA concentration was determined using the Qubit RNA High Sensitivity (HS) Kit (Invitrogen, Q32852). RNA quality and integrity were assessed using a 2100 Bioanalyzer (Agilent Technologies) with the RNA 6000 Pico Kit (Agilent, 5067-1514), and only samples with an RNA Integrity Number (RIN) > 9 were used for library preparation.

#### Library Preparation and Sequencing

RNA-seq libraries were prepared using the TruSeq Stranded Total RNA Library Prep Kit (Illumina, RS-122-2202) with 100 ng of total RNA per sample. Libraries were uniquely dual-indexed using the TruSeq UDI system (Illumina) and sequenced on a NovaSeq 6000 (Illumina) with 50 bp paired-end reads, sequenced to a depth of 20 million fragments per sample.

#### Reference genome and annotation

The human GRCh38/hg38 genome assembly was used as a reference. Gene annotation was obtained from the TxDb.Hsapiens.UCSC.hg38.knownGene Bioconductor package (https://doi.org/doi:10.18129/B9.bioc.TxDb.Hsapiens.UCSC.hg38.knownGene, version 3.15.0).

#### RNA-seq data analysis

RNA-seq experiments were sequenced paired-end with 50 bp read length. Reads were aligned to the reference genome using the STAR (version 2.7.3a)(*85*) with parameters --outFilterType BySJout, --outFilterMultimapNmax 20, --alignSJoverhangMin = 8, --alignSJDBoverhangMin 1, --outFilterMismatchNmax 999, --alignIntronMin 20, --alignIntronMax 1000000, --alignMatesGapMax 1000000, --outSAMmultNmax 1, -- outSAMtype BAM SortedByCoordinate and --outSAMunmapped Within KeepPairs. Alignments overlapping genes from the TxDb.Hsapiens.UCSC.hg38.knownGene Bioconductor package (https://doi.org/doi:10.18129/B9.bioc.TxDb.Hsapiens.UCSC.hg38.knownGene, version 3.15.0) were quantified using the qCount function with parameter orientation = “opposite” from the QuasR Bioconductor package (version 1.34.0)(*86*).

#### Differential RNA-seq analysis

To identify differentially expressed genes, lowly expressed genes were filtered out using function filterByExpr from edgeR (version 3.38.1)(*87*) using default parameters. Differentially expressed genes were identified using edgeR (version 3.38.1)(*87*). Two different models were fit for genes counts:

1. ∼ Time point (HATG-Alg 3 weeks vs. 9 weeks; HATG-Alg 3 weeks vs. 9 weeks)
2. ∼ Sample type (HATG-Alg 3 weeks vs. HATG 3 weeks; HATG-Alg 9 weeks vs. HATG 9 weeks)

For each model, dispersions are estimated using estimateDisp and statistical significance is calculated using glmQLFTest. Differentially expressed genes are defined as genes with an FDR less than 0.05.

#### Gene ontology analysis

Gene Ontology (GO) enrichment analysis was performed using the R package clusterProfiler (version 4.14) to identify biological processes (BP) significantly enriched among differentially expressed genes (DEGs). DEGs were obtained from edgeR differential expression analysis, and genes with a false discovery rate (FDR) < 0.05 were considered significant. Where indicated, DEGs were further filtered to include only genes with average logCPM > 1.

For each comparison, upregulated and downregulated gene sets were analyzed separately using the enrichGO() function, specifying the Biological Process (BP) ontology and the org.Hs.eg.db annotation package. The full set of detected genes served as the background (universe). P-values were adjusted for multiple testing using the Benjamini–Hochberg method, and redundant GO terms were reduced using the simplify() function.

### SEM

Cartilage samples were rinsed in PBS and subjected to a gradual chemical dehydration protocol involving the sequential immersion in graded ethanol solutions (30%, 50%, 70%, 90%, and 100%) for 1 h each, followed by a second series using hexamethyldisilazane (HMDS) in ethanol (30%, 50%, 70%, 90%, and 100%) for 1 h each. After the final treatment in 100% HDMS, samples were allowed to air dry in a fume hood. Once completely dried, samples were cut in half and mounted on metallic pins using carbon paste. A 10 nm carbon coating was applied using a CCU-010 Carbon Coater (Safematic) prior to imaging. Scanning electron microscopy was performed using a Zeiss Merlin microscope operated at 2 kV.

### Large Language Models (LLMs)

Large language models (ChatGPT, OpenAI) were used for language editing and proofreading during the preparation of the manuscript. No scientific content or analysis was generated using these tools.

### Statistical Analysis

All data are reported as mean ± standard deviation. Each measurement was performed on a distinct sample. Statistical analysis was performed using Python (v3.11.5) and the SciPy library (v1.11.1). Independent two-sample t-tests were performed using either the standard Student’s t-test (equal variance) or Welch’s t-test (unequal variance). Levene’s test for homogeneity of variances was used to determine equality of variances. Welch’s t-test was applied if p < 0.05, otherwise the standard Student’s t-test was used. Statistical significance was considered for a p-value below 0.05. n_d_: number of human auricular cartilage donors, n_s_: number of samples per donor.

## Acknowledgements

This work was supported by the Swiss National Science Foundation (CRSII5_173868) and Innosuisse (102.321 IP-LS). The authors gratefully acknowledge ScopeM for their support and assistance in this work. The authors further gratefully acknowledge Marine Polymer Technologies Inc. for providing sNAG nanofibrils and ViscoTec Pumpen- u. Dosiertechnik GmbH for providing the Puredyne Kit B. The authors further acknowledge the contribution of Pierre Guillon for his help with the design of the auricular 3D model and Prof. Jeffrey Bode and Dr. Sohei Majima for providing access to their laboratory’s HPLC system for the analysis of collagen crosslinks.

## Author contributions

- Conceptualization: PF, MZW
- Investigation: PF, SP, KF, AM, FR, GL, SK, APJ, DF
- Methodology: PF, SP, GL, SK
- Formal analysis: PF, SK
- Resources: FMR, DS, ENO, TL
- Supervision: PF, MZW, FMR
- Data curation: GL, SK, PF
- Visualization: PF, SK, APJ
- Writing – original draft: PF
- Writing – review & editing: all authors
- Funding acquisition: PF, MZW
- Project administration: PF, MZW

## Data and materials availability

The datasets generated and analyzed in this study are available in the ETH Zürich Research Collection repository under the doi: 10.3929/ethz-b-000737485 (*88*). All raw and processed sequencing data (RNA-seq) generated and analyzed in this study were deposited in the Gene Expression Omnibus (GEO) under GEO Series accession number GSE300497.

## References

1. D. V. Luquetti, C. L. Heike, A. V. Hing, M. L. Cunningham, T. C. Cox, Microtia: Epidemiology and genetics. American Journal of Medical Genetics Part A 158A, 124–139 (2012).

2. N. Horlock, E. Vögelin, E. T. Bradbury, A. O. Grobbelaar, D. T. Gault, Psychosocial Outcome of Patients After Ear Reconstruction: A Retrospective Study of 62 Patients. Annals of Plastic Surgery 54, (2005).

3. A. Steffen, B. Wollenberg, I. R. König, H. Frenzel, A prospective evaluation of psychosocial outcomes following ear reconstruction with rib cartilage in microtia. Journal of Plastic, Reconstructive & Aesthetic Surgery 63, 1466–1473 (2010).

4. Y. Fan, W. Liu, X. Fan, X. Niu, X. Chen, Psychosocial status of patients with unilateral and bilateral microtia before auricular reconstruction surgery. International Journal of Pediatric Otorhinolaryngology 151, 110928 (2021).

5. E. Dolgin, Advocates to bring rare disease philanthropy under one umbrella. Nature Medicine 16, 837–837 (2010).

6. I. Melnikova, Rare diseases and orphan drugs. Nature Reviews Drug Discovery 11, 267–268 (2012).

7. N. Baluch et al., Auricular reconstruction for microtia: A review of available methods. Plastic Surgery 22, 39–43 (2014).

8. F. Hartmann-Fritsch, D. Marino, E. Reichmann, About ATMPs, SOPs and GMP: The Hurdles to Produce Novel Skin Grafts for Clinical Use. Transfusion Medicine and Hemotherapy 43, 344–352 (2016).

9. D. D. Im, B. Paskhover, D. A. Staffenberg, R. Jarrahy, Current management of microtia: a national survey. Aesthetic Plast Surg 37, 402–408 (2013).

10. C. C. Breugem, K. J. Stewart, M. Kon, International Trends in the Treatment of Microtia. Journal of Craniofacial Surgery 22, (2011).

11. B. Brent, Technical Advances in Ear Reconstruction with Autogenous Rib Cartilage Grafts: Personal Experience with 1200 Cases. Plast Reconstr Surg 104, (1999).

12. Y. Kawanabe, S. Nagata, A New Method of Costal Cartilage Harvest for Total Auricular Reconstruction: Part I. Avoidance and Prevention of Intraoperative and Postoperative Complications and Problems. Plast Reconstr Surg 117, (2006).

13. Y. Kawanabe, S. Nagata, A New Method of Costal Cartilage Harvest for Total Auricular Reconstruction: Part II. Evaluation and Analysis of the Regenerated Costal Cartilage. Plast Reconstr Surg 119, (2007).

14. F. Firmin, State-of-the-Art Autogenous Ear Reconstruction in Cases of Microtia. Advances in Oto-Rhino-Laryngology 68, 25–52 (2010).

15. Z. Sun et al., Costal Cartilage Assessment in Surgical Timing of Microtia Reconstruction. Journal of Craniofacial Surgery 28, (2017).

16. A. Yamada, Autologous Rib Microtia Construction: Nagata Technique. Facial Plast Surg Clin North Am 26, 41–55 (2018).

17. A. L. Johns, R. E. Lucash, D. D. Im, S. L. Lewin, Pre and post-operative psychological functioning in younger and older children with microtia. Journal of Plastic, Reconstructive & Aesthetic Surgery 68, 492–497 (2015).

18. X. Long, N. Yu, J. Huang, X. Wang, Complication rate of autologous cartilage microtia reconstruction: a systematic review. Plast Reconstr Surg Glob Open 1, e57 (2013).

19. Y. Y. Fu, C. L. Li, J. L. Zhang, T. Y. Zhang, Autologous cartilage microtia reconstruction: Complications and risk factors. Int J Pediatr Otorhinolaryngol 116, 1–6 (2019).

20. J. F. Reinisch, Y. Tahiri, in Modern Microtia Reconstruction: Art, Science, and New Clinical Techniques, J. F. Reinisch, Y. Tahiri, Eds. (Springer International Publishing, Cham, 2019), pp. 91–110.

21. J. F. Reinisch, in Modern Microtia Reconstruction: Art, Science, and New Clinical Techniques, J. F. Reinisch, Y. Tahiri, Eds. (Springer International Publishing, Cham, 2019), pp. 321–334.

22. R. K. Sharma et al., Early Postoperative Complications in Microtia Reconstruction: An Analysis of the NSQIP-P Database. The Laryngoscope 134, 1214–1219 (2024).

23. S. Cugno, N. Bulstrode, in Modern Microtia Reconstruction: Art, Science, and New Clinical Techniques, J. F. Reinisch, Y. Tahiri, Eds. (Springer International Publishing, Cham, 2019), pp. 135–141.

24. P. A. Federspil, Auricular Prostheses in Microtia. Facial Plastic Surgery Clinics 26, 97–104 (2018).

25. A. S. Murthy, in Modern Microtia Reconstruction: Art, Science, and New Clinical Techniques, J. F. Reinisch, Y. Tahiri, Eds. (Springer International Publishing, Cham, 2019), pp. 291–302.

26. A. J. Lin, J. L. Bernstein, J. A. Spector, Ear Reconstruction and 3D Printing: Is It Reality? Current Surgery Reports 6, 4 (2018).

27. R. Roy et al., Analysis of bending behavior of native and engineered auricular and costal cartilage. Journal of Biomedical Materials Research Part A 68A, 597–602 (2004).

28. C. A. Vacanti, L. G. Cima, D. Ratkowski, J. Upton, J. P. Vacanti, Tissue Engineered Growth of New Cartilage in the Shape of a Human Ear Using Synthetic Polymers Seeded with Chondrocytes. MRS Proceedings 252, 367 (1991).

29. A. He et al., Cell yield, chondrogenic potential, and regenerated cartilage type of chondrocytes derived from ear, nasoseptal, and costal cartilage. Journal of Tissue Engineering and Regenerative Medicine 12, 1123–1132 (2018).

30. M. F. b. Ishak et al., The formation of human auricular cartilage from microtic tissue: An in vivo study. International Journal of Pediatric Otorhinolaryngology 79, 1634–1639 (2015).

31. B. P. Cohen et al., Long-Term Morphological and Microarchitectural Stability of Tissue-Engineered, Patient-Specific Auricles In Vivo. Tissue Engineering Part A 22, 461–468 (2016).

32. X. Dong et al., Three-Dimensional-Printed External Scaffolds Mitigate Loss of Volume and Topography in Engineered Elastic Cartilage Constructs. CARTILAGE 13, 1780S–1789S (2021).

33. X. Dong et al., Efficient engineering of human auricular cartilage through mesenchymal stem cell chaperoning. Journal of Tissue Engineering and Regenerative Medicine, (2022).

34. N. Hirano et al., Ethanol treatment of nanoPGA/PCL composite scaffolds enhances human chondrocyte development in the cellular microenvironment of tissue-engineered auricle constructs. PLOS ONE 16, e0253149 (2021).

35. Z. Yin et al., Regeneration of elastic cartilage with accurate human-ear shape based on PCL strengthened biodegradable scaffold and expanded microtia chondrocytes. Applied Materials Today 20, 100724 (2020).

36. G. Zhou et al., In Vitro Regeneration of Patient-specific Ear-shaped Cartilage and Its First Clinical Application for Auricular Reconstruction. EBioMedicine 28, 287–302 (2018).

37. B. P. Cohen, J. L. Bernstein, K. A. Morrison, J. A. Spector, L. J. Bonassar, Tissue engineering the human auricle by auricular chondrocyte-mesenchymal stem cell co-implantation. PLOS ONE 13, e0202356 (2018).

38. P. Fisch, N. Broguiere, S. Finkielsztein, T. Linder, M. Zenobi-Wong, Bioprinting of Cartilaginous Auricular Constructs Utilizing an Enzymatically Crosslinkable Bioink. Advanced Functional Materials 31, 2008261 (2021).

39. D. Zielinska et al., Combining bioengineered human skin with bioprinted cartilage for ear reconstruction. Sci. Adv. 9, eadh1890 (2023).

40. D. Gvaramia et al., Evaluation of Bioprinted Autologous Cartilage Grafts in an Immunocompetent Rabbit Model. Adv. Ther. 7, (2024).

41. Y. Yao, C. Wang, Dedifferentiation: inspiration for devising engineering strategies for regenerative medicine. npj Regenerative Medicine 5, 14 (2020).

42. M. Gosset, F. Berenbaum, S. Thirion, C. Jacques, Primary culture and phenotyping of murine chondrocytes. Nature Protocols 3, 1253–1260 (2008).

43. E. W. Mandl et al., Fibroblast growth factor-2 in serum-free medium is a potent mitogen and reduces dedifferentiation of human ear chondrocytes in monolayer culture. Matrix Biology 23, 231–241 (2004).

44. E. W. Mandl, S. W. van der Veen, J. A. Verhaar, G. J. van Osch, Serum-free medium supplemented with high-concentration FGF2 for cell expansion culture of human ear chondrocytes promotes redifferentiation capacity. Tissue Eng 8, 573–580 (2002).

45. Y. Xie et al., FGF/FGFR signaling in health and disease. Signal Transduction and Targeted Therapy 5, 181 (2020).

46. I. Martin, G. Vunjak-Novakovic, J. Yang, R. Langer, L. E. Freed, Mammalian Chondrocytes Expanded in the Presence of Fibroblast Growth Factor 2 Maintain the Ability to Differentiate and Regenerate Three-Dimensional Cartilaginous Tissue. Experimental Cell Research 253, 681–688 (1999).

47. N. G. M. Thielen, P. M. van der Kraan, A. P. M. van Caam, TGFβ/BMP Signaling Pathway in Cartilage Homeostasis. Cells 8, (2019).

48. P. M. van der Kraan, The changing role of TGFβ in healthy, ageing and osteoarthritic joints. Nature Reviews Rheumatology 13, 155–163 (2017).

49. I. Martin et al., Enhanced cartilage tissue engineering by sequential exposure of chondrocytes to FGF-2 during 2D expansion and BMP-2 during 3D cultivation. Journal of Cellular Biochemistry 83, 121–128 (2001).

50. P. Fisch, N. Broguiere, S. Finkielsztein, T. Linder, M. Zenobi-Wong. (ETH Zurich Research Collection, 2021).

51. G.-Z. Jin, H.-W. Kim, Efficacy of collagen and alginate hydrogels for the prevention of rat chondrocyte dedifferentiation. Journal of Tissue Engineering 9, 2041731418802438 (2018).

52. H.-p. Lee, L. Gu, D. J. Mooney, M. E. Levenston, O. Chaudhuri, Mechanical confinement regulates cartilage matrix formation by chondrocytes. Nature Materials 16, 1243–1251 (2017).

53. N. Broguiere, E. Cavalli, G. M. Salzmann, L. A. Applegate, M. Zenobi-Wong, Factor XIII Cross-Linked Hyaluronan Hydrogels for Cartilage Tissue Engineering. ACS Biomater Sci Eng 2, 2176–2184 (2016).

54. D. Gvaramia, J. Kern, Y. Jakob, M. Zenobi-Wong, N. Rotter, Regenerative Potential of Perichondrium: A Tissue Engineering Perspective. Tissue Eng Part B Rev, (2021).

55. E. A. Makris, D. J. Responte, N. K. Paschos, J. C. Hu, K. A. Athanasiou, Developing functional musculoskeletal tissues through hypoxia and lysyl oxidase-induced collagen cross-linking. Proc. Natl. Acad. Sci. 111, E4832 (2014).

56. C. E. H. Schmelzer, L. Duca, Elastic fibers: formation, function, and fate during aging and disease. The FEBS Journal n/a, (2021).

57. A. Sorushanova et al., The Collagen Suprafamily: From Biosynthesis to Advanced Biomaterial Development. Advanced Materials 31, 1801651 (2019).

58. J. K. Lee et al., Tension stimulation drives tissue formation in scaffold-free systems. Nature materials 16, 864–873 (2017).

59. A. Heinz, Elastic fibers during aging and disease. Ageing Research Reviews 66, 101255 (2021).

60. N. Miekus et al., MMP-14 degrades tropoelastin and elastin. Biochimie 165, 32–39 (2019).

61. S. A. Hibbert, R. E. B. Watson, C. E. M. Griffiths, N. K. Gibbs, M. J. Sherratt, Selective proteolysis by matrix metalloproteinases of photo-oxidised dermal extracellular matrix proteins. Cellular Signalling 54, 191–199 (2019).

62. R. J. Nims et al., Matrix Production in Large Engineered Cartilage Constructs Is Enhanced by Nutrient Channels and Excess Media Supply. Tissue Engineering Part C: Methods 21, 747–757 (2014).

63. A. D. Cigan et al., Nutrient channels and stirring enhanced the composition and stiffness of large cartilage constructs. Journal of Biomechanics 47, 3847–3854 (2014).

64. Michael B. Albro et al., Accumulation of Exogenous Activated TGF-β in the Superficial Zone of Articular Cartilage. Biophysical Journal 104, 1794–1804 (2013).

65. I. Pomerantseva et al., Ear-Shaped Stable Auricular Cartilage Engineered from Extensively Expanded Chondrocytes in an Immunocompetent Experimental Animal Model. Tissue Eng Part A 22, 197–207 (2016).

66. Y. Liu et al., Double network hydrogels encapsulating genetically modified dedifferentiated chondrocytes for auricular cartilage regeneration. J. Mater. Chem. B 13, 1823–1844 (2024).

67. C. E. H. Schmelzer, L. Duca, Elastic fibers: formation, function, and fate during aging and disease. The FEBS Journal 289, 3704–3730 (2022).

68. N. F. Tojais et al., Codependence of Bone Morphogenetic Protein Receptor 2 and Transforming Growth Factor-β in Elastic Fiber Assembly and Its Perturbation in Pulmonary Arterial Hypertension. Arteriosclerosis Thrombosis Vasc Biology 37, 1559–1569 (2018).

69. D. Hubmacher, D. P. Reinhardt, in The Extracellular Matrix: an Overview, R. P. Mecham, Ed. (Springer Berlin Heidelberg, Berlin, Heidelberg, 2011), pp. 233–265.

70. K. Tiedemann, B. Bätge, P. K. Müller, D. P. Reinhardt, Interactions of Fibrillin-1 with Heparin/Heparan Sulfate, Implications for Microfibrillar Assembly *. Journal of Biological Chemistry 276, 36035–36042 (2001).

71. L. Sabatier et al., Heparin/heparan sulfate controls fibrillin-1, −2 and −3 self-interactions in microfibril assembly. FEBS Letters 588, 2890–2897 (2014).

72. D. J. Responte, B. Arzi, R. M. Natoli, J. C. Hu, K. A. Athanasiou, Mechanisms underlying the synergistic enhancement of self-assembled neocartilage treated with chondroitinase-ABC and TGF-β1. Biomaterials 33, 3187–3194 (2012).

73. J. H. Haga, Y.-S. J. Li, S. Chien, Molecular basis of the effects of mechanical stretch on vascular smooth muscle cells. Journal of Biomechanics 40, 947–960 (2007).

74. L. Venkataraman, C. A. Bashur, A. Ramamurthi, Impact of Cyclic Stretch on Induced Elastogenesis Within Collagenous Conduits. Tissue Engineering Part A 20, 1403–1415 (2013).

75. C. A. Nietupski, A. Moset Zupan, S. C. Schutte, Impact of Cyclic Strain on Elastin Synthesis in a 3D Human Myometrial Culture Model. Tissue Engineering Part C: Methods 30, 279–288 (2024).

76. R. S. Allen, S. K. Biswas, A. W. Seifert, Ear pinna growth and differentiation is conserved in murids and requires BMP signaling for chondrocyte proliferation. Dev. (Camb., Engl.) 152, DEV204560 (2025).

77. R. Yang et al., BMP signaling maintains auricular chondrocyte identity and prevents microtia development by inhibiting protein kinase A. eLife 12, RP91883 (2024).

78. M. Minoux et al., Mouse Hoxa2 mutations provide a model for microtia and auricle duplication. Development 140, 4386–4397 (2013).

79. N. Si et al., Identification of loss-of-function HOXA2 mutations in Chinese families with dominant bilateral microtia. Gene 757, 144945 (2020).

80. T. C. Cox, E. D. Camci, S. Vora, D. V. Luquetti, E. E. Turner, The genetics of auricular development and malformation: New findings in model systems driving future directions for microtia research. European Journal of Medical Genetics 57, 394–401 (2014).

81. D. A. Bichara et al., Successful Creation of Tissue-Engineered Autologous Auricular Cartilage in an Immunocompetent Large Animal Model. Tissue Engineering Part A 20, 303–312 (2013).

82. N. Broguiere, L. Isenmann, M. Zenobi-Wong, Novel enzymatically cross-linked hyaluronan hydrogels support the formation of 3D neuronal networks. Biomaterials 99, 47–55 (2016).

83. T. F. Linsenmayer, M. J. C. Hendrix, Monoclonal antibodies to connective tissue macromolecules: Type II collagen. Biochemical and Biophysical Research Communications 92, 440–446 (1980).

84. J. Perła-Kaján, A. Gryszczyńska, S. Mielcarek, H. Jakubowski, Cation exchange HPLC analysis of desmosines in elastin hydrolysates. Analytical and Bioanalytical Chemistry 401, 2473–2479 (2011).

85. A. Dobin et al., STAR: ultrafast universal RNA-seq aligner. Bioinformatics 29, 15–21 (2013).

86. D. Gaidatzis, A. Lerch, F. Hahne, M. B. Stadler, QuasR: quantification and annotation of short reads in R. Bioinformatics 31, 1130–1132 (2015).

87. M. D. Robinson, D. J. McCarthy, G. K. Smyth, edgeR: a Bioconductor package for differential expression analysis of digital gene expression data. Bioinformatics 26, 139–140 (2010).

88. P. Fisch et al. (ETH Zurich).

