## Supplementary Figures for "Tissue Engineered Elastic Cartilage-Mimetic Auricular Grafts for Ear Reconstruction"

### Supplementary Material

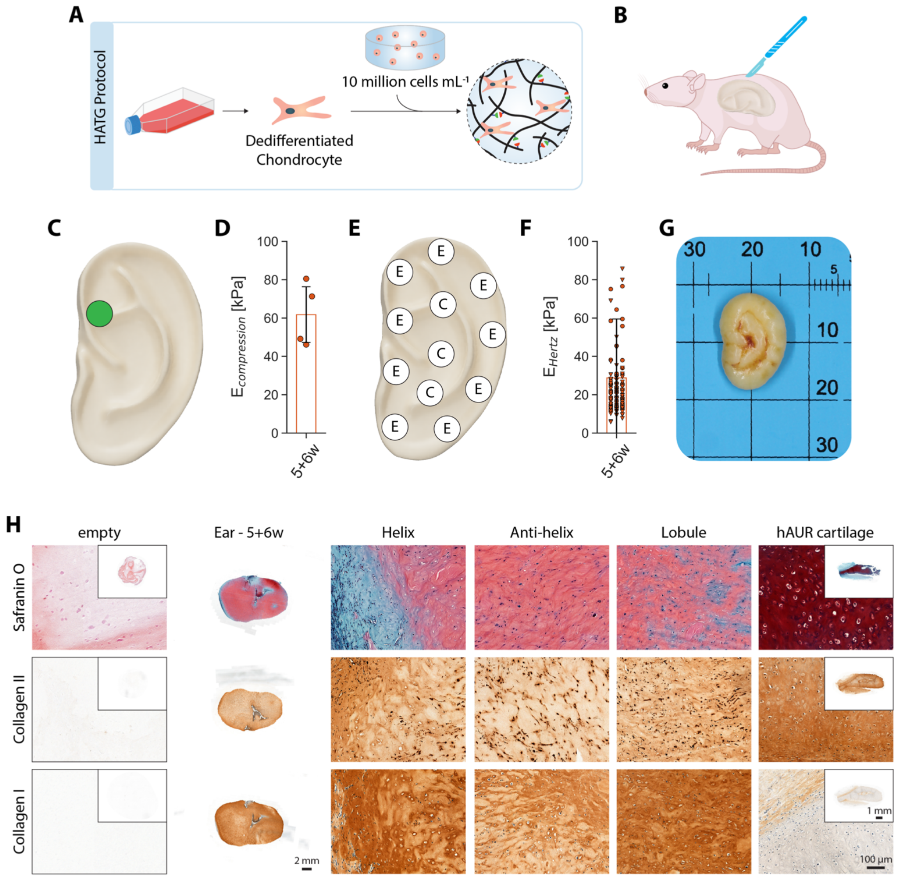


**Fig. S1 | Subcutaneous implantation of auricular grafts consisting of fibrocartilage in rats – HATG protocol.** (**A,B**) Illustration of the implantation in rats. (**C**) Position of the ear graft which was used for compression tests. (**D**) Compressive modulus of ears. (**E**) Positions of the ear graft where indentations tests were carried out (E: edge, C: center). (**F**) Hertz modulus obtained from indentation tests along the edge (▾) and in the center (•) of the ear. (**G**) Picture of a collapsed ear after 6 weeks *in vivo*. (**H**) Histology and immunohistochemistry for glycosaminoglycans (Safranin O), collagen II and collagen I of ears after maturing for 5 weeks *in vitro* and 6 weeks *in vivo*, showing the deposition of fibrocartilage. Empty HATG bioink and human auricular (hAUR) cartilage as control. Scale bar: ears: 2 mm, controls: full view: 1 mm, zoom in: 100 µm. $n_d = 1$, $n_s = 4$.

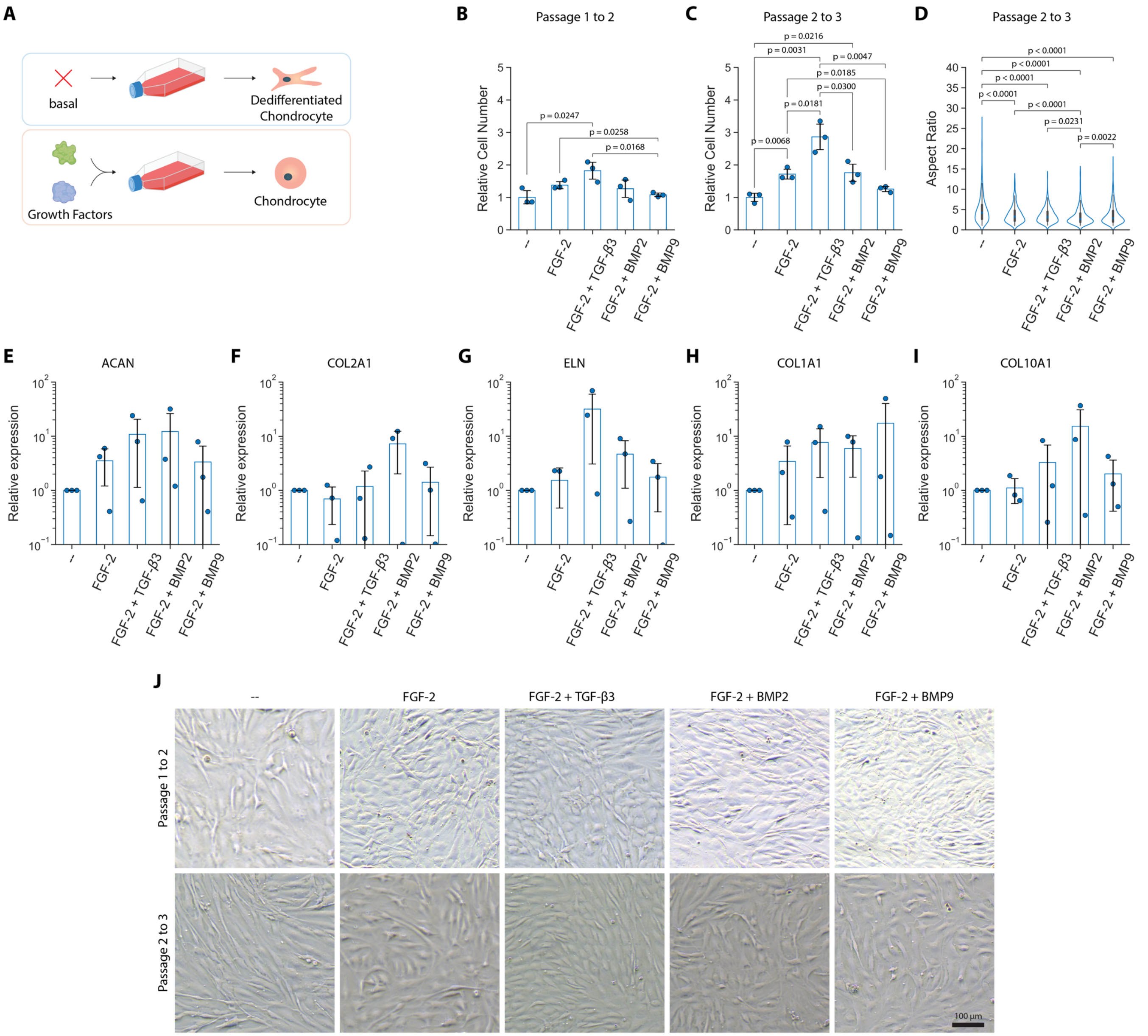


**Fig. S2 | Growth factor supplementation during hAUR expansion.** (**A**), Illustration of the supplementation of growth factors during 2D expansion of hAUR. (**B-C**), Relative cell number compared to the basal condition when cells were expanded with various growth factors at concentrations of 10 ng $mL^{-1}$ per growth factor from passage 1 to passage 2 (**B**) and passage 2 to passage 3 (**C**). (**D**), Aspect ratio of hAUR expanded in the presence of various growth factors as shown in (**J**). (**E-I**), Relative gene expression compared to basal conditions (DMEM 31966, 10% v/v FBS and 10 µg $mL^{-1}$ gentamicin) of hAUR expanded in the presence of various growth factors (10 ng $mL^{-1}$) for ACAN, elastin, collagen II, collagen I and collagen X based on $2^{-\Delta\Delta Ct}$ at passage 3. (**J**), Images of hAUR expanded in the presence of various growth factors. Scale bar: 100 µm. $n_d$ = 3, $n_s$ = 1.

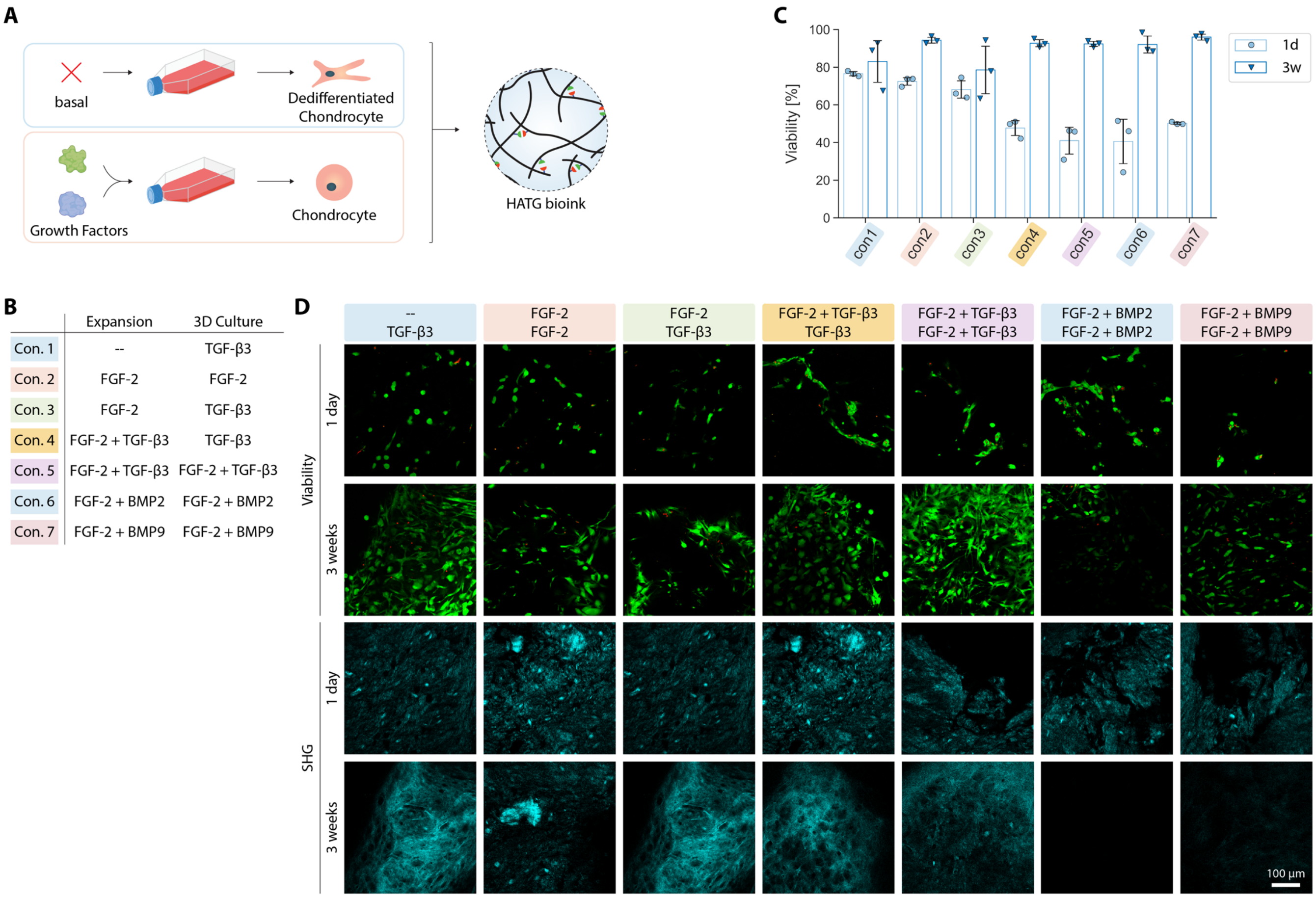


**Fig. S3 | Growth factor supplementation during hAUR expansion and subsequent encapsulation in the HATG bioink at a density of 10 million cells per mL.** (**A**) Illustration of the supplementation of growth factors during 2D expansion of hAUR and their combination with the HATG bioink at 10 million cells per mL. (**B**) Overview of the conditions (Con.) tested. (**C**) Viability of hAUR encapsulated in the HATG bioink after 1 day and 3 weeks. (**D**) Representative viability images (live: green, dead: red) of hAUR in the HATG bioink after 1 day and 3 weeks as well as second harmonic generation (SHG) images. Scale bar: 100 µm. $n_d$ = 1, $n_s$ = 3.

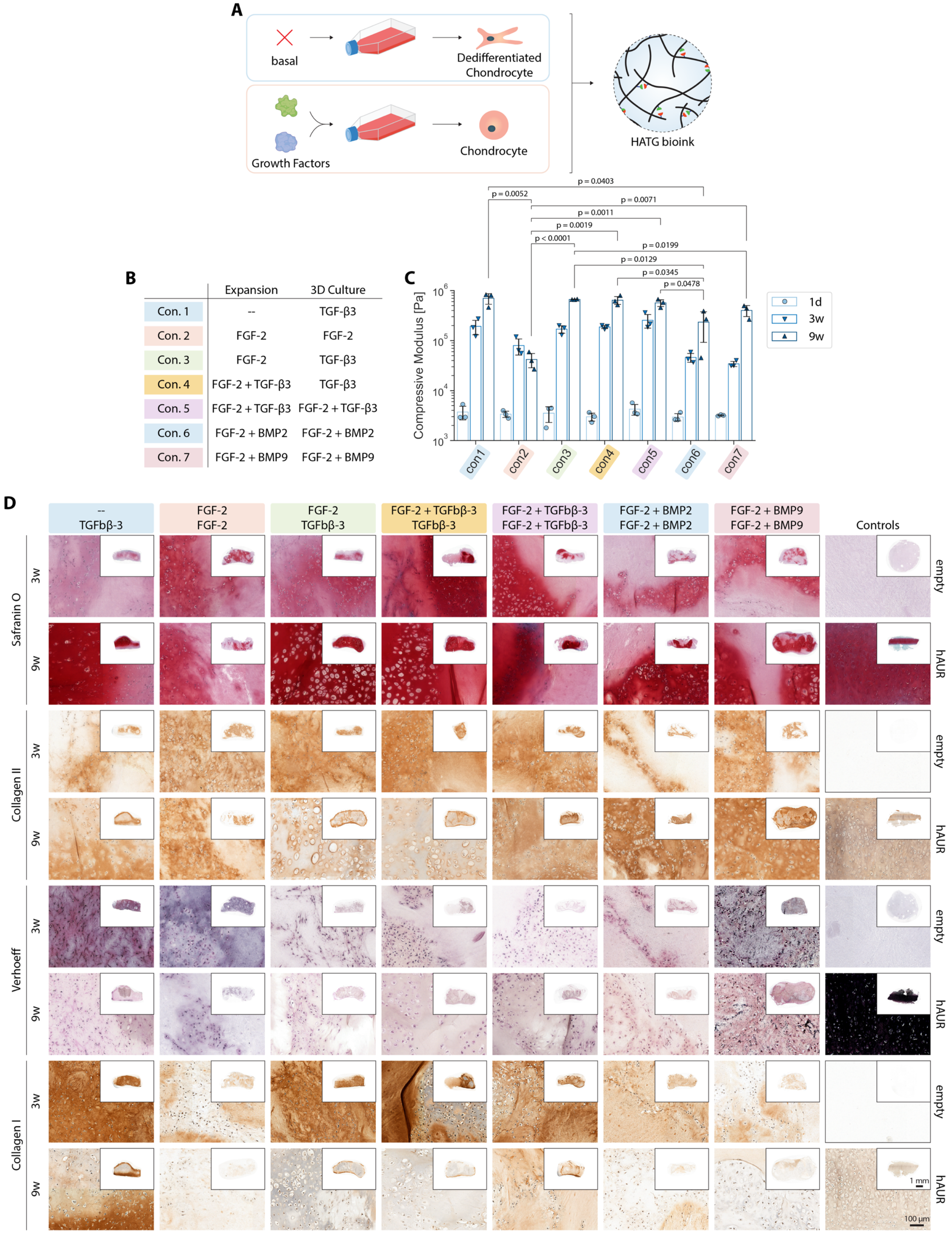


**Fig. S4 | Growth factor supplementation during hAUR expansion and subsequent encapsulation in the HATG bioink at a density of 10 million cells per mL - continued.** (**A**), Illustration of the supplementation of growth factors during 2D expansion of hAUR and their combination with the HATG bioink at 10 million cells per mL. (**B**), Overview of the conditions (Con.) tested. (**C**), Compressive modulus of grafts after 1 day and , 3 and 9 weeks. (**D**), Histology of grafts after 3 and 9 weeks compared to empty bioink gels and human auricular (hAUR) cartilage as control. Scale bar: close up: 100 µm, inserts: 1 mm. $n_d = 1$, $n_s = 3$.

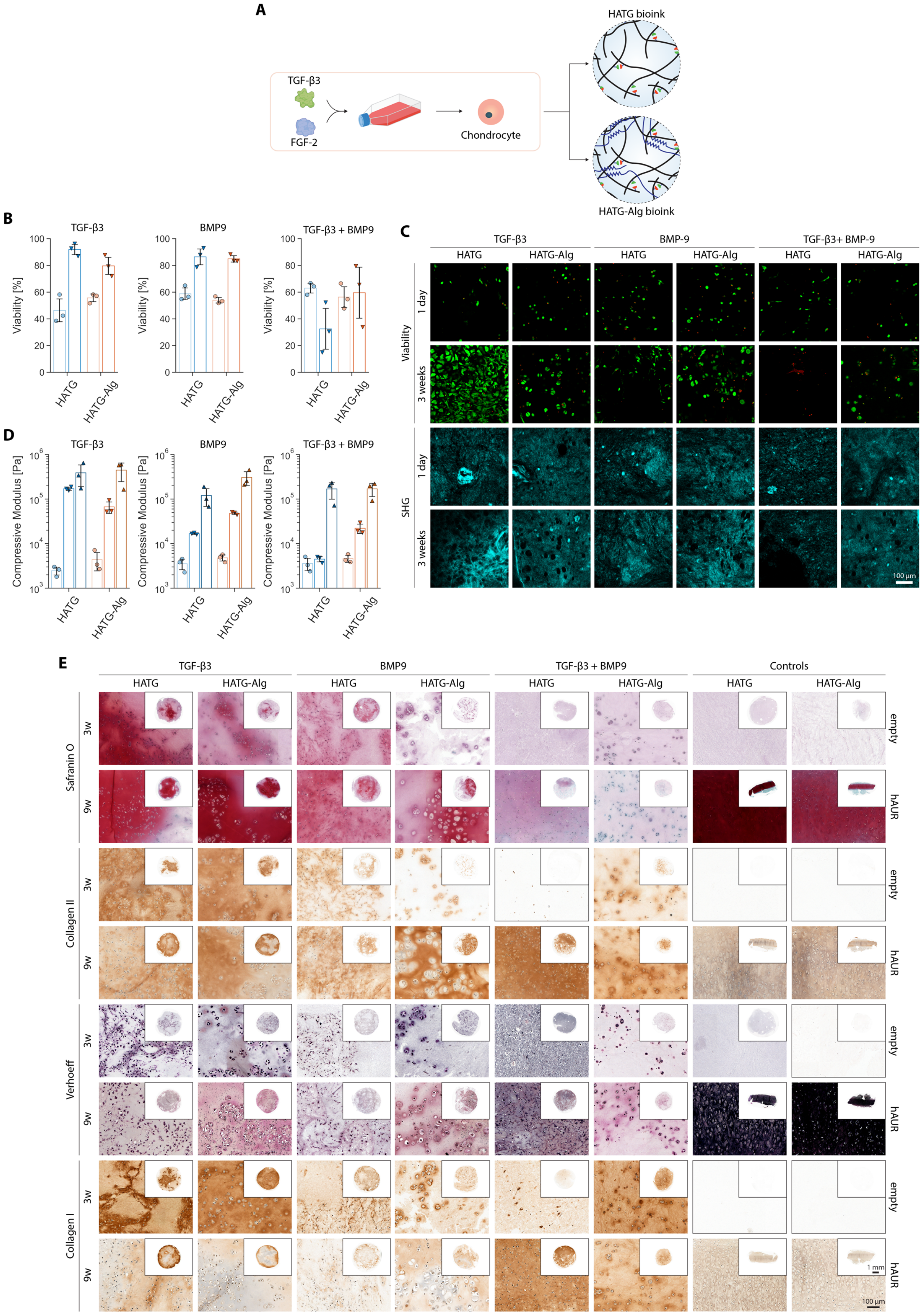
A
TGF-β3
FGF-2
Chondrocyte
HATG bioink
HATG-Alg bioink
B
TGF-β3
BMP9
TGF-β3 + BMP9
Viability [%]
HATG
HATG-Alg
C
TGF-β3
BMP-9
TGF-β3+ BMP-9
HATG
HATG-Alg
Viability
SHG
1 day
3 weeks
100 µm
D
Compressive Modulus [Pa]
E
Controls
Safranin O
Collagen II
Verhoeff
Collagen I
3w
9w
empty
hAUR
1 mm
100 µm

**Fig. S5 | Growth factor supplementation during hAUR expansion and subsequent encapsulation in the either the HATG or HATG-Alg bioink at a density of 10 million cells per mL.** hAUR were expanded in the presence of FGF-2 and TGF-β3 and grafts were cultured in the presence of TGF-β3, BMP-9 or TGF-β3 and BMP-9. (**A**) Illustration of the supplementation of growth factors during 2D expansion of hAUR and their combination with either the HATG or HATG-Alg bioink at 10 million cells per mL. (**B**) Viability of hAUR encapsulated in either bioink after 1 day (•) and 3 weeks ( ▲ ). (**C**) Representative viability images (live: green, dead: red) of hAUR after 1 day and 3 weeks as well as second harmonic generation (SHG) images. Scale bar: 100 µm. (**D**) Compressive modulus of grafts after 1 day ( • ) and 3 ( ▼ ) and 9 ( ▲ ) weeks. (**E**) Histology of grafts after 3 and 9 weeks compared to empty bioink gels and human auricular (hAUR) cartilage as control. Scale bar: close up: 100 µm, inserts: 1 mm. $n_d = 1$, $n_s = 3$.

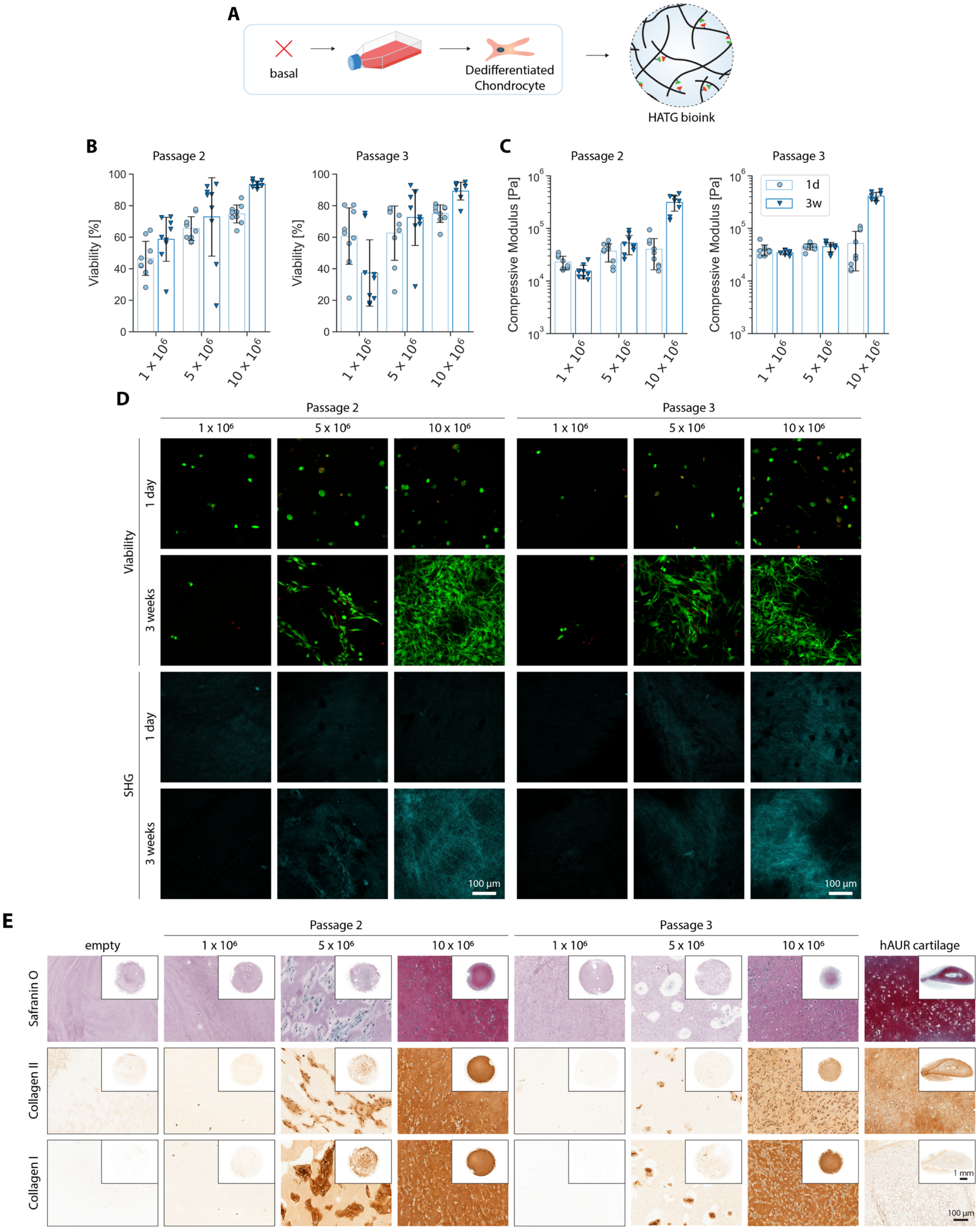


**Fig. S6 | Impaired maturation of HATG bioink grafts at a lower cell density of 1, 5 and 10 million cells per mL after passage 2 and 3.** (**A**) hAUR were expanded in the absence of additional growth factors and combined with the HATG bioink at 1, 5 and 10 million cells per mL. (**B**) Viability of hAUR encapsulated in either bioink after 1 day (•) and 3 weeks ( ▴ ). (**C**) Compressive modulus of grafts after 1 day (•) and 3 weeks ( ▾ ). (**D**) Representative viability images (live: green, dead: red) of hAUR after 1 day and 3 weeks as well as second harmonic generation (SHG) images. Scale bar: 100 µm. (**E**) Histology of grafts after 3 weeks compared to empty bioink gels and human auricular (hAUR) cartilage as control. Scale bar: close up: 100 µm, inserts: 1 mm. $n_d = 3$, $n_s = 3$.

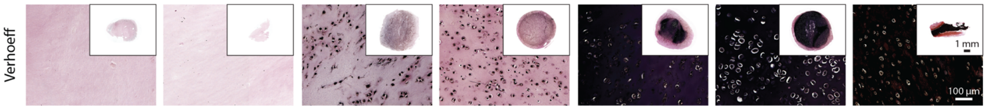


**Fig. S7 |** Original, unadjusted Verhoeff staining compared to empty bioink gels and human auricular (hAUR). Scale bar: close up: 100 µm, inserts: 1 mm. $n_d$ = 1, $n_s$ = 3.

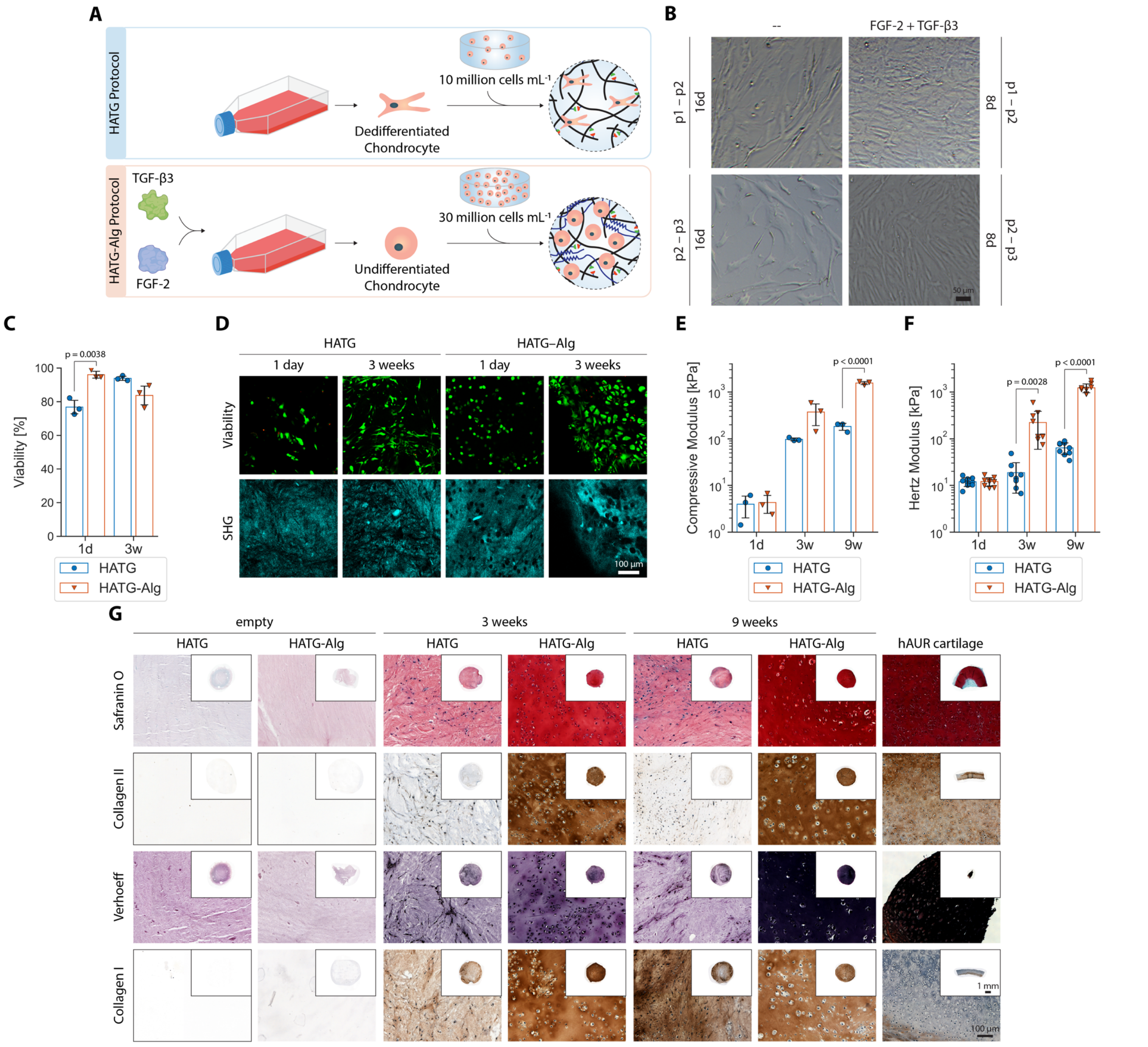


**Fig. S8 | Transcriptome analysis of grafts matured using the HATG and HATG-Alg protocol – part 1.** (**A**) HATG Protocol: hAUR were expanded in the absence of additional growth factors and combined with the HATG bioink at a density of 10 million cells per mL. HATG-Alg Protocol: hAUR were expanded in the presence of 10 ng $mL^{-1}$ FGF-2 and TGF-β3 and combined with the HATG-Alg bioink at a density of 30 million cells per mL. (**B**) Brightfield images of hAUR expanded either in basal media (--) or basal media supplemented with 10 ng $mL^{-1}$ FGF-2 and TGF-β3 (FGF-2 + TGF-β3) taken when cells were passaged (passage 1 to 2 and passage 2 to 3). Without the addition of growth factors during expansion, cells required 16 days to reach confluency, whereas they only required 8 days in the presence of growth factors. Scale bar: 50 µm. (**C**) Cell viability in the HATG and HATG-Alg bioink 1 day and 3 weeks after printing. (**D**) Representative images of cell viability (green = viable, red = dead) and second harmonic generation for either bioink after 1 day and 3 weeks. Scale bar: 100 µm. (**E**) Compressive and (**F**) Hertz modulus of grafts after 1 day and 3 and 9 weeks. (**G**) Histology of grafts after 3 and 9 weeks compared to empty bioink gels and human auricular (hAUR) cartilage as control. Scale bar: close up: 100 µm, inserts: 1 mm. $n_d = 1$, $n_s = 3$.

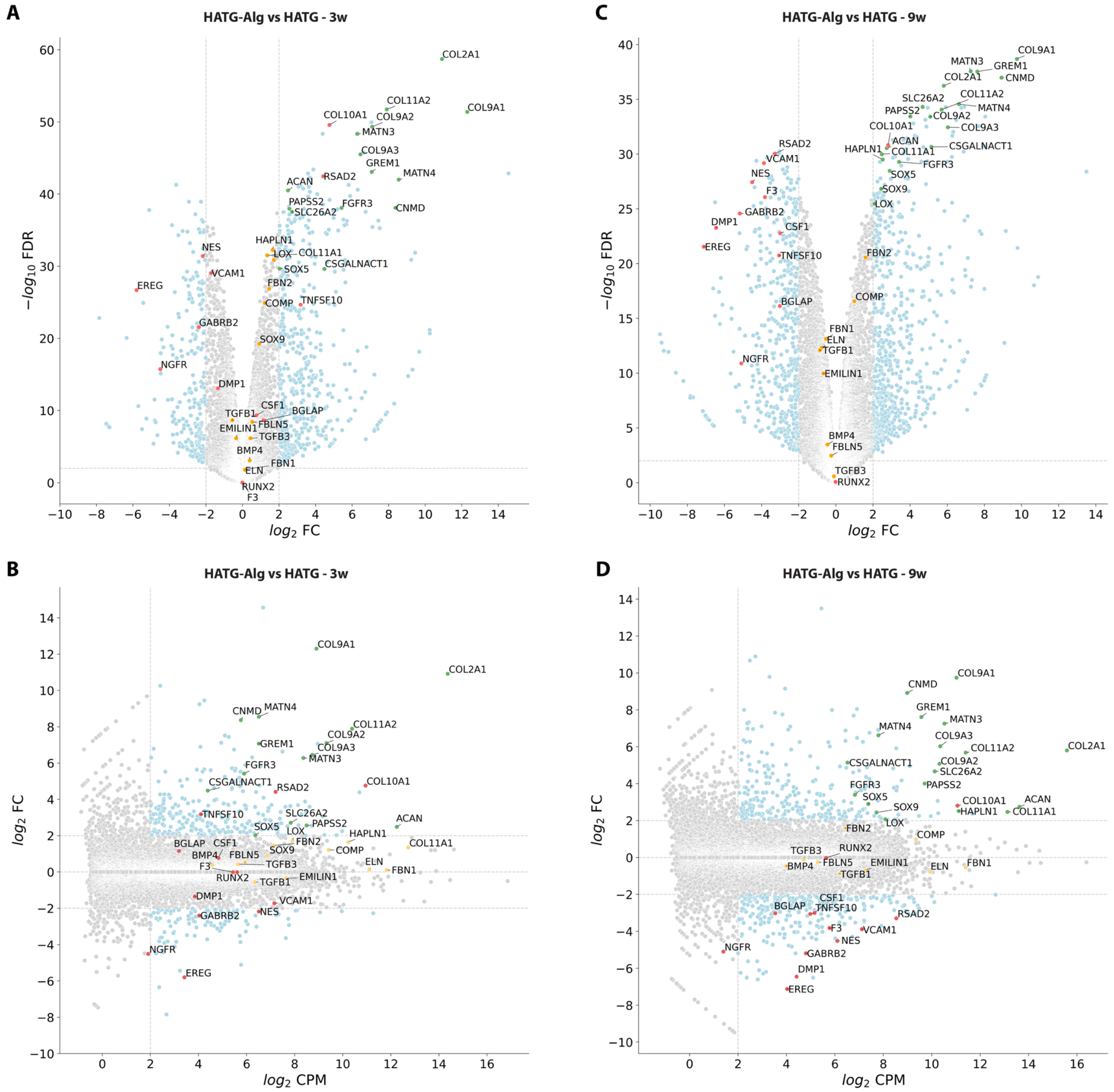


**Fig. S9 | Transcriptome analysis of grafts matured using the HATG and HATG-Alg protocol – part 2.** Volcano (top) and MA (bottom) plots comparing samples of the different protocols and timepoints: HATG-Alg compared to the HATG protocol at 3 weeks (**A-B**, HATG-Alg vs. HATG - 3w) and 9 weeks (**C-D**, HATG-Alg vs. HATG - 3w). Green: genes positively influencing cartilage formation, red: genes negatively influencing cartilage formation and/or involved in other pathways, orange: further genes related to elastic cartilage formation.

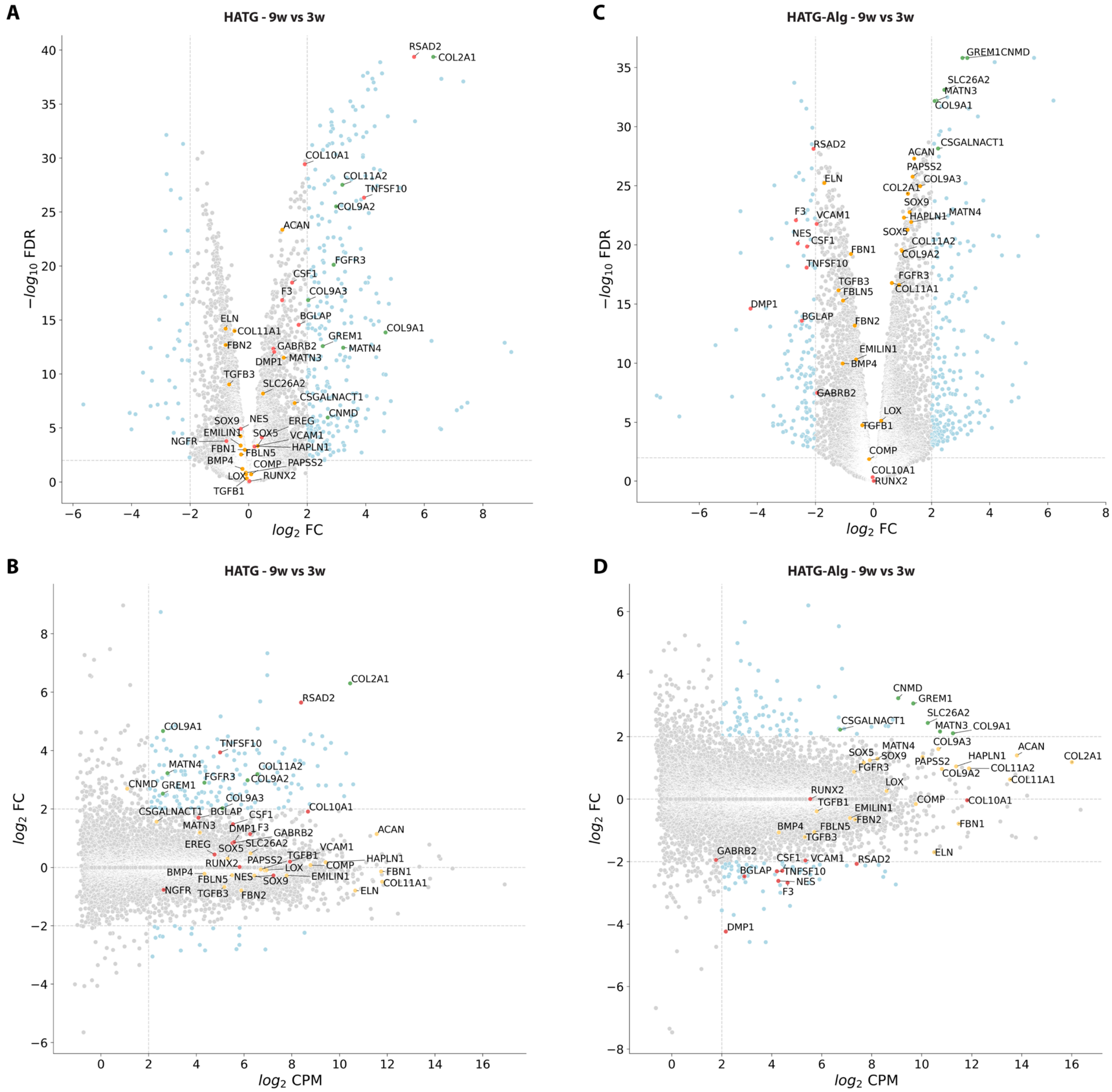


**Fig. S10 | Transcriptome analysis of grafts matured using the HATG and HATG-Alg protocol – part 3.** Volcano (top) and MA (bottom) plots comparing samples of the different protocols and timepoints: 9 weeks compared to 3 weeks for the HATG (**A-B**, HATG - 9w vs. 3w) and HATG-Alg (**C-D**, HATG-Alg - 9w vs. 3w) protocol. Green: genes positively influencing cartilage formation, red: genes negatively influencing cartilage formation and/or involved in other pathways, orange: further genes related to elastic cartilage formation.

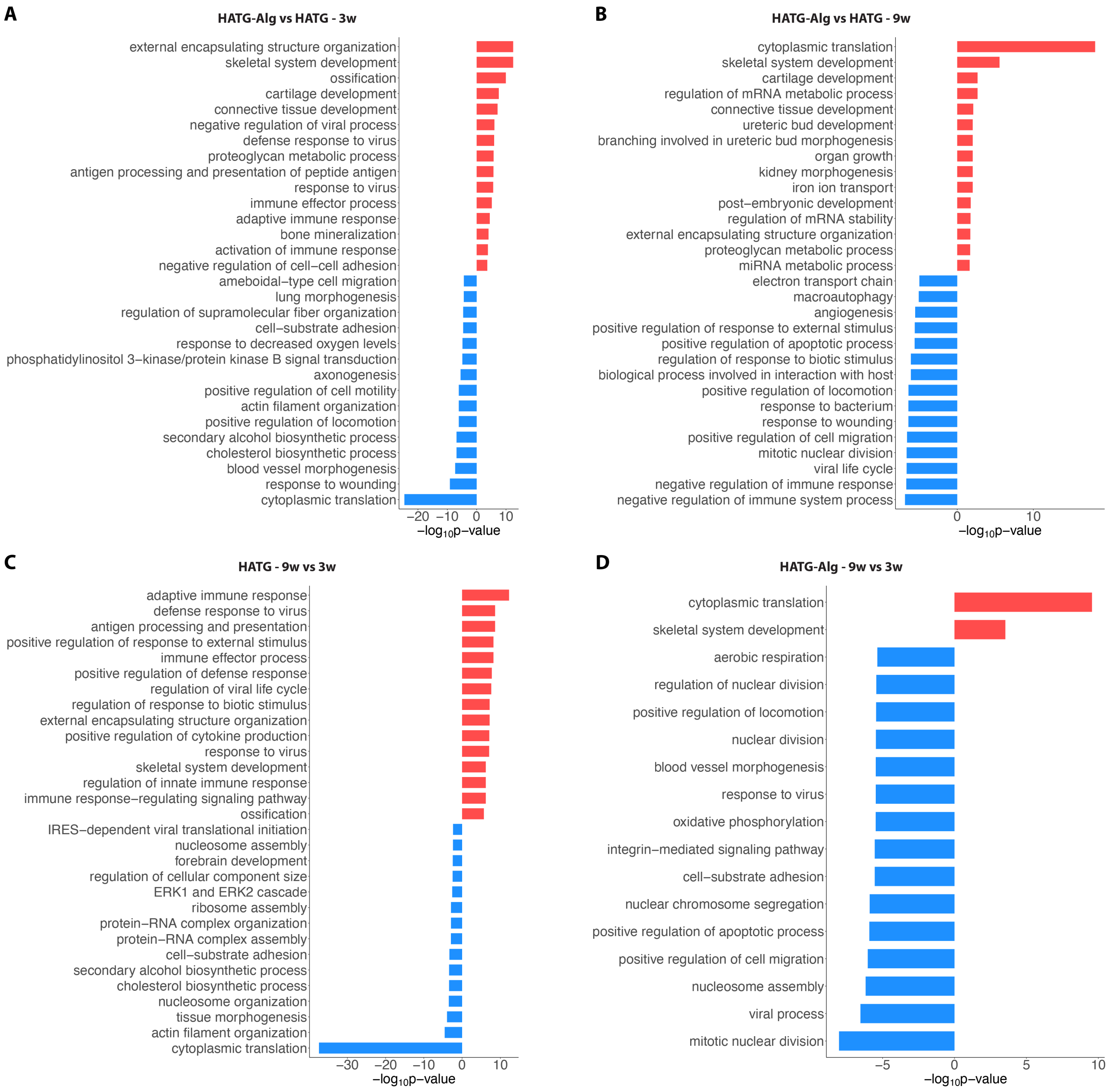


**Fig. S11 | Transcriptome analysis of grafts matured using the HATG and HATG-Alg protocol – part 4.** Gene ontology (GO) analysis of samples of the different protocols and timepoints: HATG-Alg compared to the HATG protocol at 3 weeks (**A**, HATG-Alg vs. HATG - 3w, red: upregulated in HATG-Alg, blue: downregulated in HATG-Alg) and 9 weeks (**B**, HATG-Alg vs. HATG - 3w, red: upregulated in HATG-Alg, blue: downregulated in HATG-Alg) and 9 weeks compared to 3 weeks for the HATG (**C**, HATG - 9w vs. 3w, red: upregulated at 9 weeks, blue: downregulated at 9 weeks) and HATG-Alg (**D**, HATG-Alg - 9w vs. 3w, red: upregulated at 9 weeks, blue downregulated at 9 week) protocol (average logCPM > 1). Top 15 GO:BP terms.

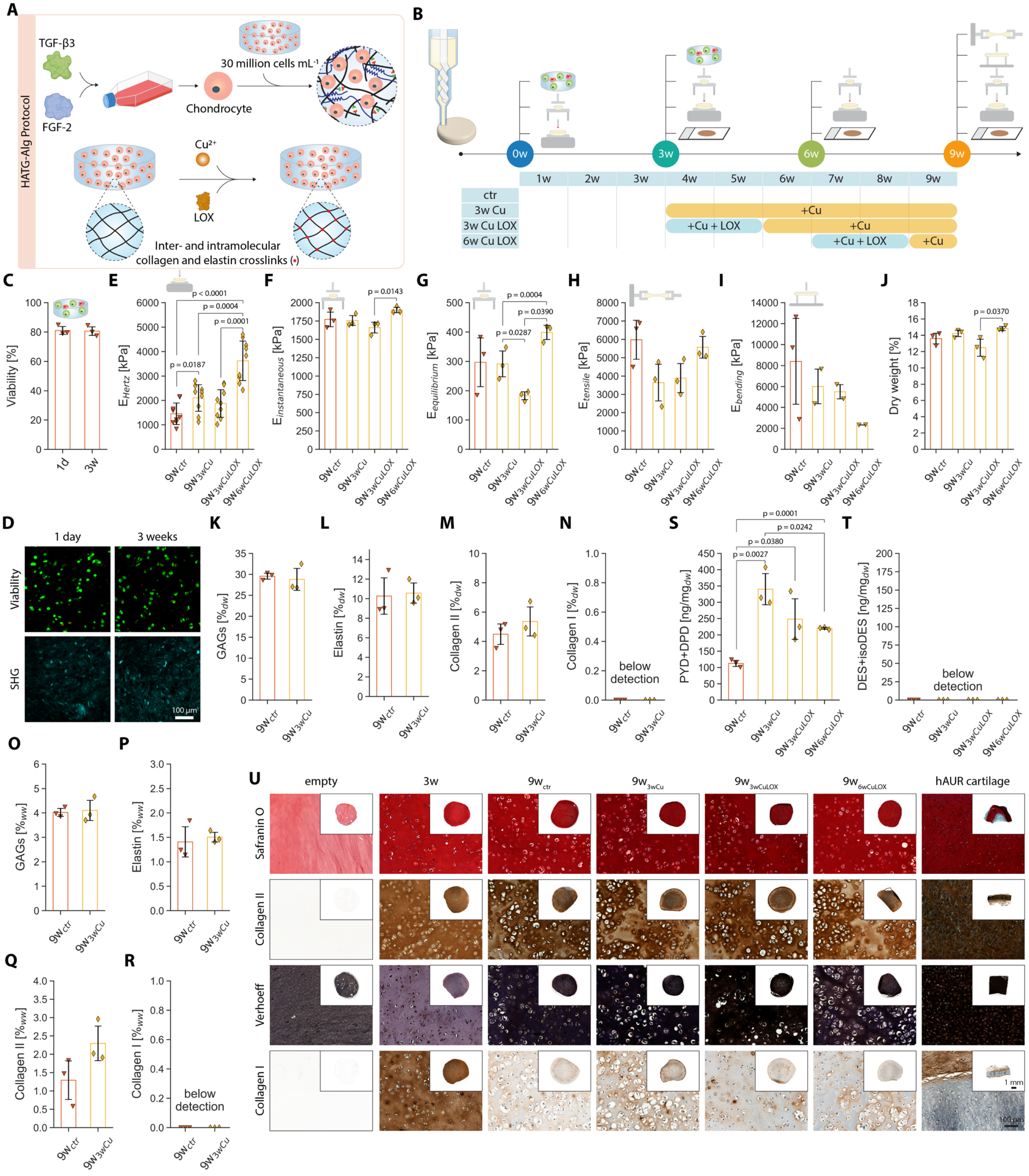


**Fig. S12 | Supplementation of culture media with copper and/or exogenous lysyl oxidase.** (**A**) Grafts were prepared following the HATG-Alg protocol. In addition, copper and/or lysyl oxidase (LOX) were added to the graft culture media to enhance inter- and intra-molecular crosslinks in collagen and elastin. (**B**) Timeline of the supplementation regime. Copper (+Cu) was added after maturing grafts for 3 weeks and then maintained for the remaining 6 weeks until grafts reached 9 weeks (3w Cu). In addition, two conditions in which Cu and LOX were supplemented (+Cu +LOX) were tested. Cu and LOX were supplemented for 2 weeks either after the 3rd week of culture (3w Cu LOX) or after the 6th week of culture (6w Cu LOX). Afterwards, culture media was supplemented with Cu until grafts reached 9 weeks. Grafts without Cu or LOX supplementation were run as control (ctr). (**C**) Cell viability in the HATG and HATG-Alg bioink 1 day and 3 weeks after printing. (**D**) Representative images of cell viability (green = viable, red = dead) and second harmonic generation for either bioink after 1 day and 3 weeks. Scale bar: 100 µm. (**E-I**) Mechanical characterization of grafts after 9 weeks of maturation in indentation (**E**), compression stress-relaxation (**F,G**), tension (**H**) and bending (**I**). (**J**), Dry weight of samples. (**K-N**) GAGs, elastin, collagen II and collagen I as percentage dry weight ($\%_{dw}$). (**O-R**) GAGs elastin, collagen II and collagen I as percentage wet weight ($\%_{ww}$). (**S**) pyridinoline and deoxypyridinoline and (**T**) desmosine and isodesmosine crosslinks in collagen and elastin respectively. (**U**) Histology of grafts after 3 and 9 weeks compared to empty bioink gels and human auricular (hAUR) cartilage as control. Scale bar: close up: 100 µm, inserts: 1 mm. $n_d = 1$, $n_s = 3$.

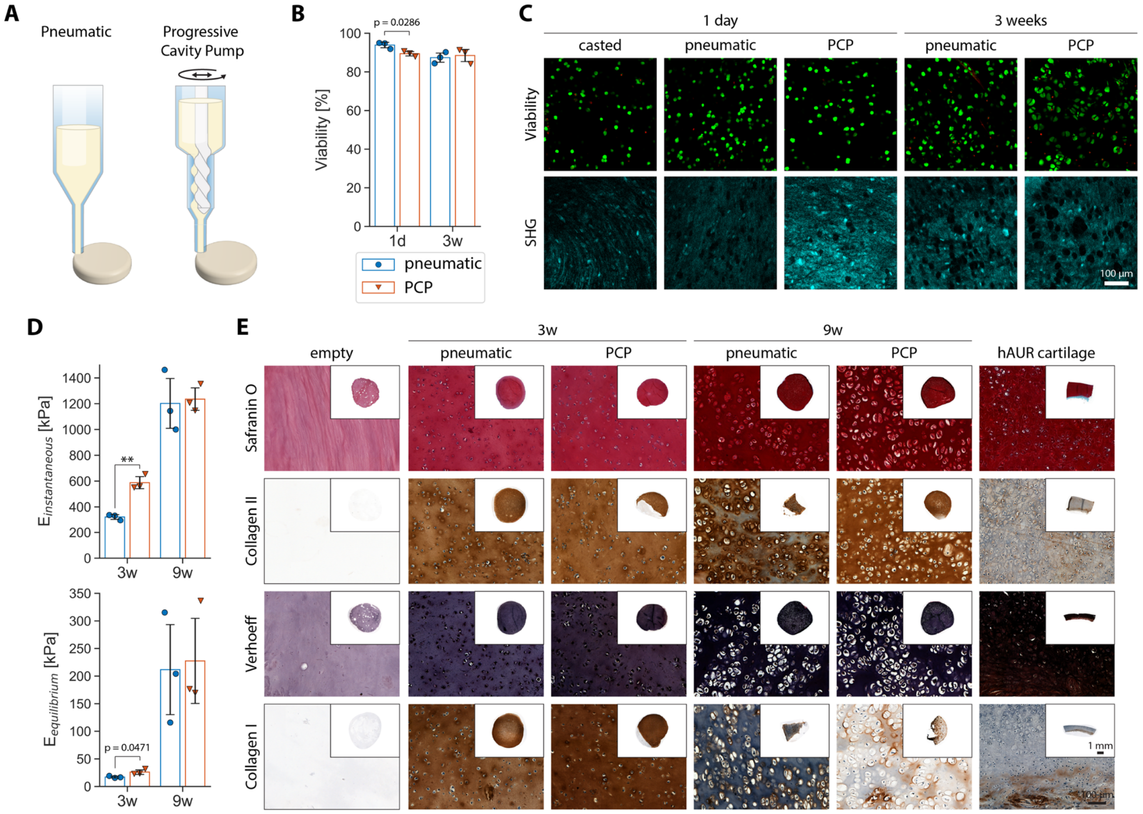


**Fig. S13 | Progressive cavity pump-based bioprinting.** (**A**) Illustration of the working principle of a progressive cavity pump (PCP) compared to the traditionally used pneumatic extrusion. (**B**) Cell viability of grafts printed with pneumatic and PCP-based extrusion 1 day and 3 weeks after printing. (**C**) Representative images of cell viability (green = viable, red = dead) and second harmonic generation for either bioink after 1 day and 3 weeks. Scale bar: 100 µm. (**D**) Instantaneous ($E_{instantaneous}$) and equilibrium ($E_{equilibrium}$) modulus of grafts after 3 and 9 weeks printed with pneumatic and PCP-based extrusion. (**E**) Histology of grafts after 3 and 9 weeks compared to empty bioink gels and human auricular (hAUR) cartilage as control. Scale bar: close up: 100 µm, inserts: 1 mm. $n_d = 1$, $n_s = 3$.

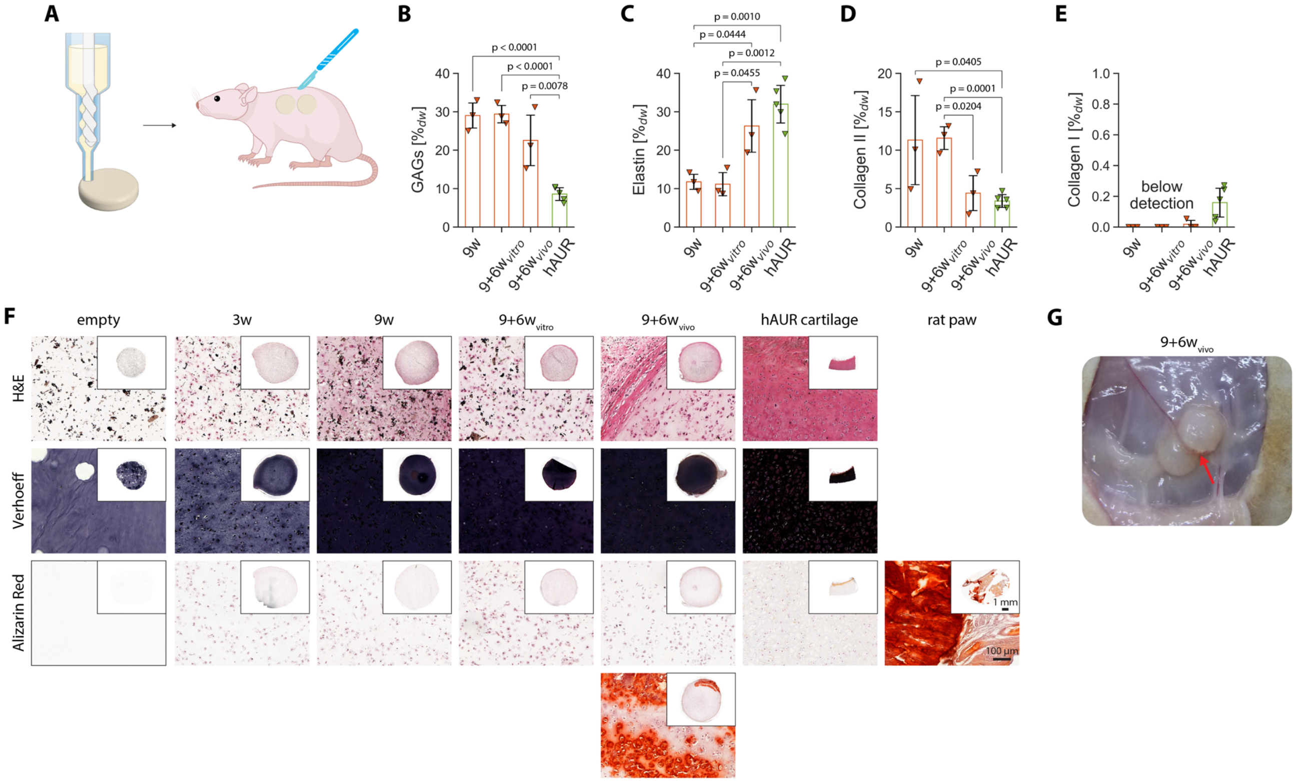


**Fig. S14 | *In vivo* implantation of tissue engineered elastic cartilage grafts – cylinders.** (**A**) Grafts were bioprinted using a progressive cavity pump-based extruder, matured for 9 weeks and implanted subcutaneously in immunocompromised rats for 6 weeks. (**B-E**) Evaluation of graft ECM components: GAGs, elastin, collagen II and collagen I per dry weight ($\%_{dw}$). (**F**) Histological evaluation of tissue engineered elastic cartilage compared to human auricular cartilage, rat paw and empty HATG-Alg samples. Original, unadjusted Verhoeff staining. (**G**) Image of grafts after 6 weeks *in vivo* with one graft showing slight calcification. 9w: 9-week culture, 9+6w vitro: 9+6-week culture, 9+6w vivo: 9-week culture plus 6 weeks *in vivo*. $n_d = 1$, $n_s = 3$.

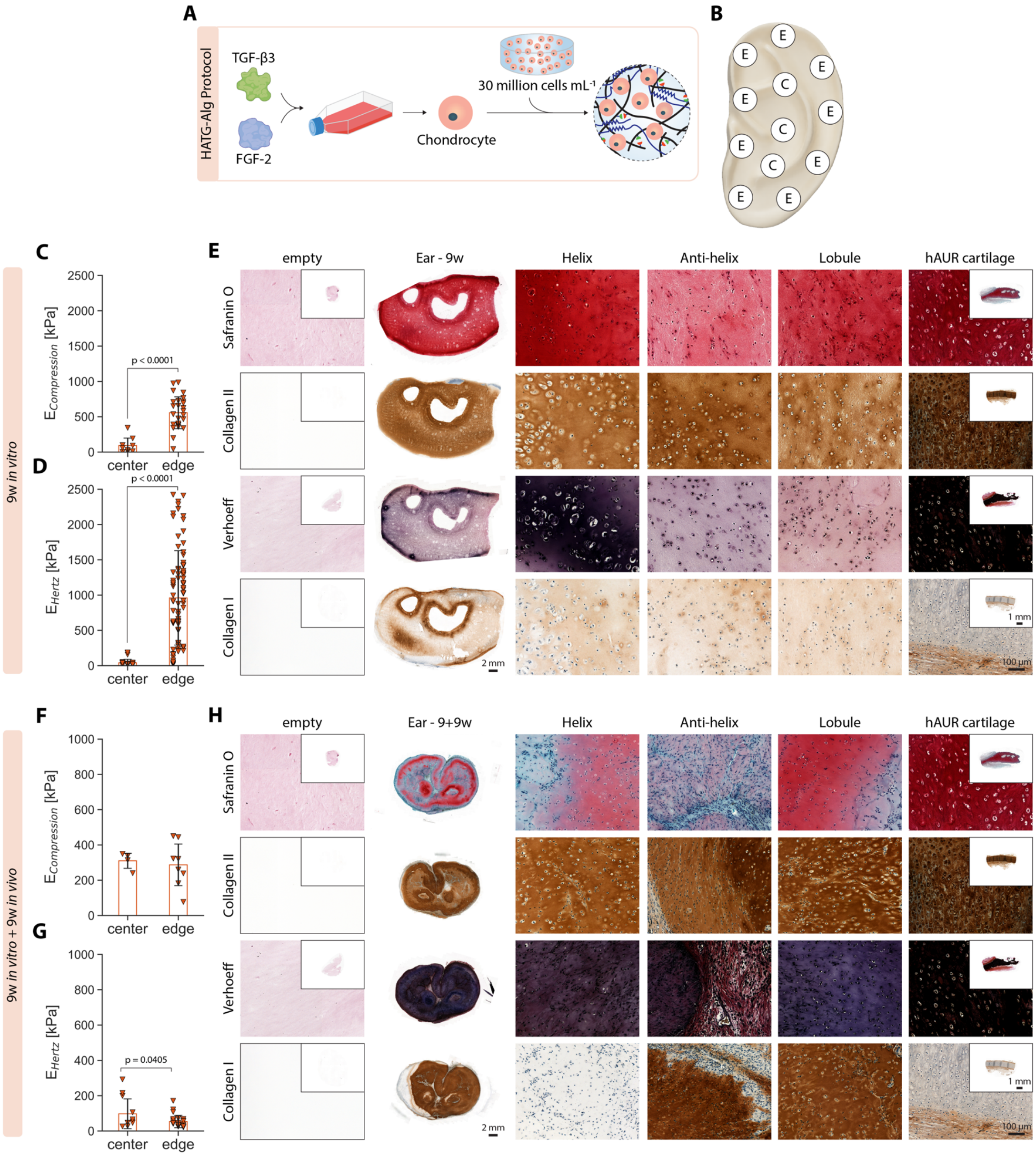


**Fig. S15 | Inhomogeneous maturation of bioprinted auricular grafts – in vitro and in vivo.** (**A**) Combination of growth factor expansion of hAUR to prevent their dedifferentiation, addition of alginate to increase stress relaxation of the bioink and increase of cell density from 10 to 30 million cells per mL to counter insufficient maturation of grafts. (**B**) Positions of the ear graft where compression tests were carried out (E: edge, C: center). (**C-D**) Compressive ($E_{compression}$) and Hertz ($E_{Hertz}$) modulus along the edge and in the center of auricular grafts after 9 weeks *in vitro*. (**E**) Histology of auricular grafts after 9 weeks *in vitro* compared to empty bioink gels and human auricular (hAUR) cartilage as control. Scale bar: close up: 100 µm, inserts: 1 mm. $n_d$ = 1, $n_s$ = 3. (**F-G**) Compressive ($E_{compression}$) and Hertz ($E_{Hertz}$) modulus along the edge and in the center of auricular grafts after 9 weeks *in vitro* and an additional 9 weeks *in vivo*. (**H**) Histology of auricular grafts after 9 weeks maturation *in vitro* and an additional 9 weeks subcutaneous *in vivo* compared to empty bioink gels and human auricular (hAUR) cartilage as control. Scale bar: close up: 100 µm, inserts: 1 mm. $n_d$ = 1, $n_s$ = 4.

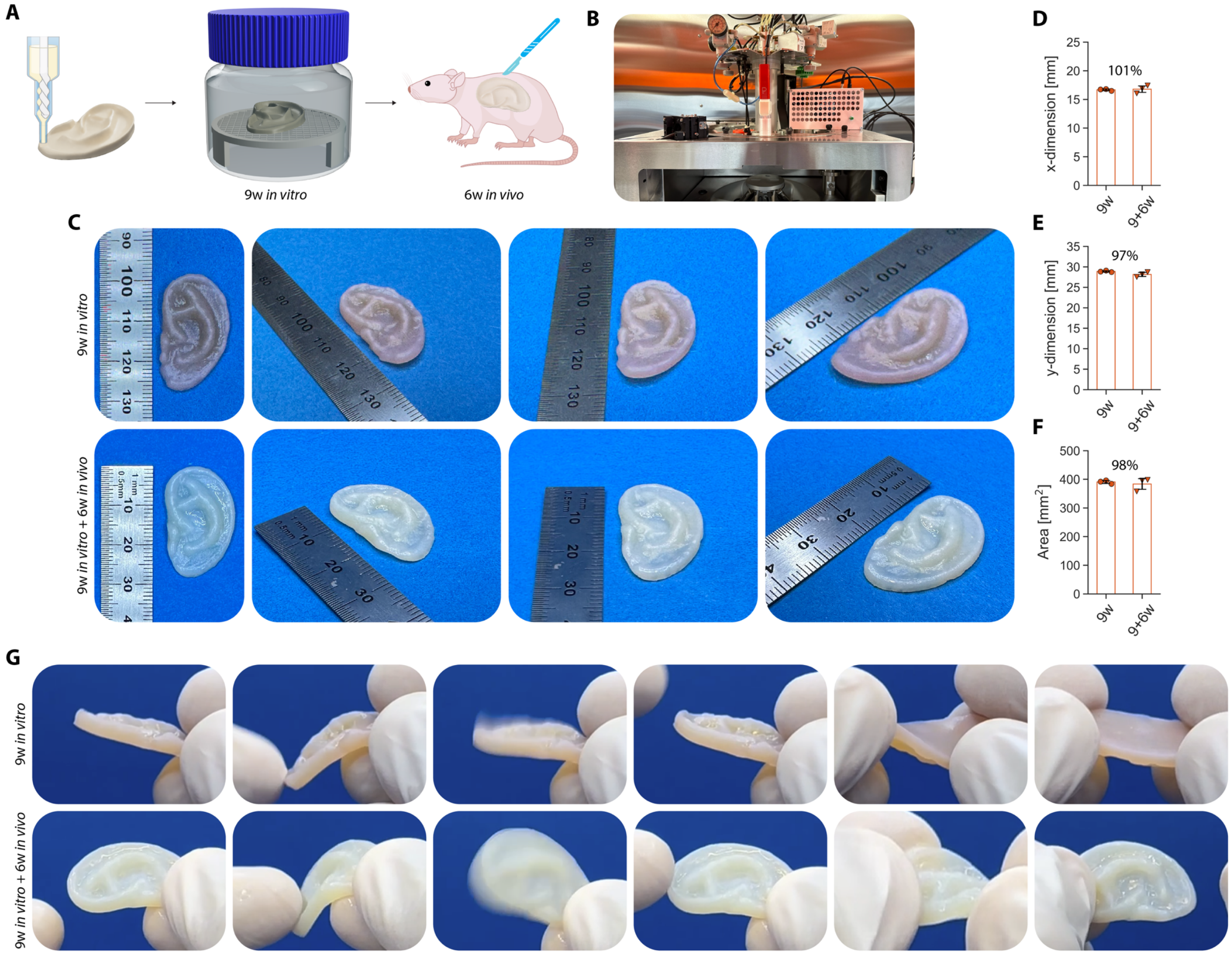


**Fig. S16 | *In vivo* evaluation of tissue engineered elastic cartilage-mimetic auricular grafts.** (**A**) Auricular grafts were bioprinted using a PCP-based extruder, cultured on a custom designed bioreactor platform to allow nutrient perfusion from all sides of the graft for 9 weeks, and implanted subcutaneously in immunocompromised rats for 6 weeks. (**B**) Image of the bioprinting setup with the PCP-based extruder (Puredyne Kit) installed in the Biofactory (regenHU). (**C**) Images of auricular grafts after 9 weeks of *in vitro* maturation (9w *in vitro*) and an additional 6 weeks *in vivo* (9w *in vitro* + 6w *in vivo*). (**D-F**) Quantification of the width (x-dimension), length (y-dimension) and area of auricular grafts after 9 weeks *in vitro* and an additional 6 weeks *in vivo*. (**G**) Image sequence of auricular grafts being deformed and regaining their original shape (Movie S1). $n_d = 1$, $n_s = 3$.

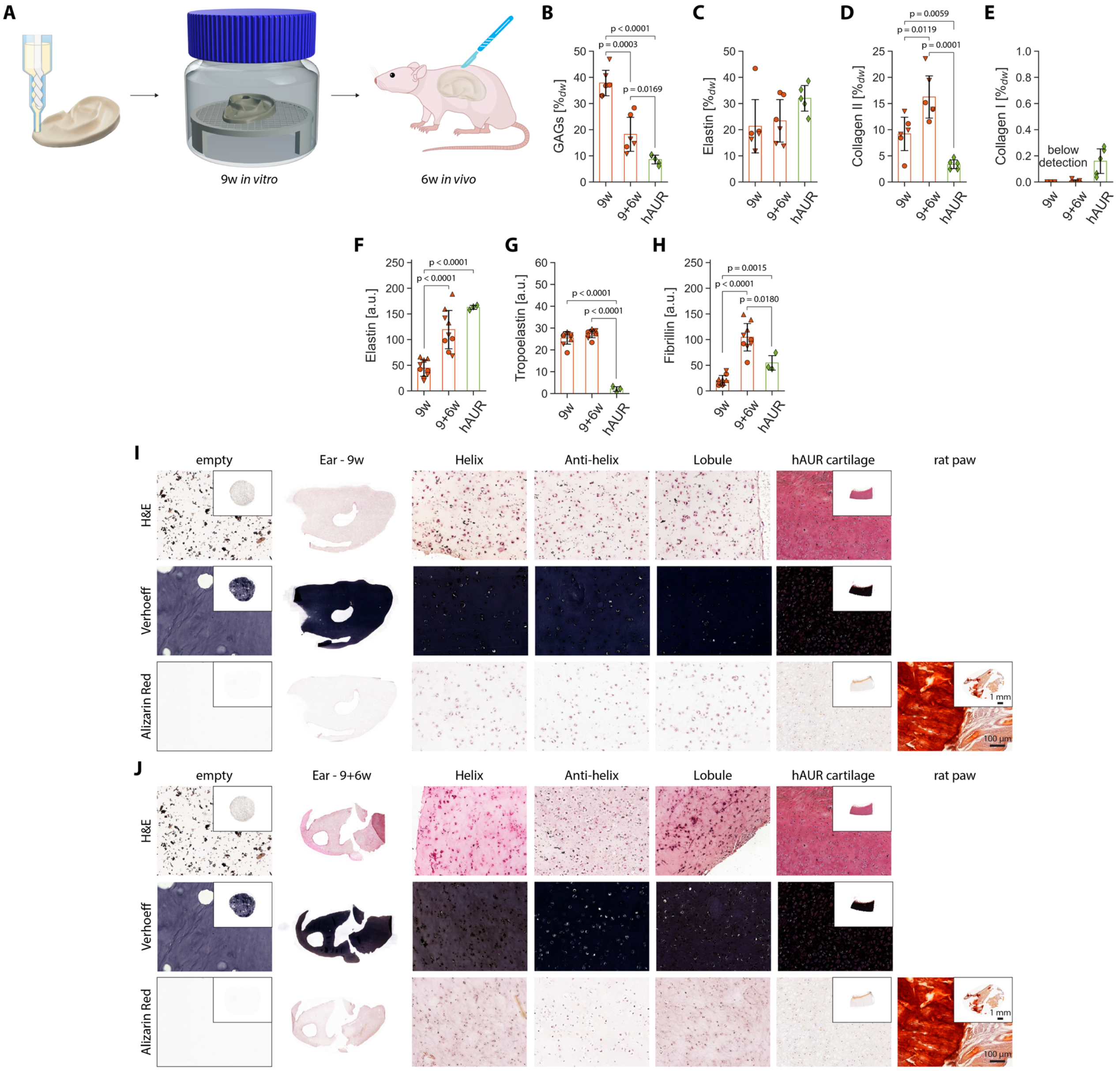


**Fig. S17 | *In vivo* evaluation of tissue engineered elastic cartilage-mimetic auricular grafts.** (**A**) Auricular grafts were bioprinted using a PCP-based extruder, cultured on a custom designed bioreactor platform to allow nutrient perfusion from all sides of the graft for 9 weeks, and implanted subcutaneously in immunocompromised rats for 6 weeks. (**B-E**) Evaluation of graft ECM components: GAGs, elastin, collagen II and collagen I per dry weight ($\%_{dw}$). (**F-H**) Color deconvolution of immunohistochemical stainings for elastin, tropoelastin and fibrillin-1. (**I-J**) Histological stainings for hematoxylin and eosin (H&E) and Alizarin Red compared to empty bioink gels, human auricular (hAUR) cartilage and rat paw as control. Original, unadjusted Verhoeff staining. Scale bar: close up: 100 µm, inserts: 1 mm. $n_d = 1$, $n_s = 3$.
